# Inflation induced motility for long-distance vertical migration

**DOI:** 10.1101/2022.08.19.504465

**Authors:** Adam G. Larson, Rahul Chajwa, Hongquan Li, Manu Prakash

## Abstract

The daily vertical migrations of plankton play a crucial role in shaping marine ecosystems and influencing global biogeochemical cycles. They also form the foundation of the largest daily biomass movement on Earth. Surprisingly, amongst this diverse group of organisms, some single cell protists transit these depths exceeding 50 meters without employing flagella or cilia, and the underlying mechanisms remain poorly understood. It has been previously proposed that this capability relies on the cell’s ability to regulate its internal density relative to seawater. Here, using *Pyrocystis noctiluca* as a model system, we demonstrate the primary mechanism for this density control is a rapid cellular inflation event, during which a single plankton cell expands its volume six-fold in less than 10 minutes. This self-regulated cellular inflation selectively imports fluid less dense than surrounding seawater, and can effectively sling-shot a cell and reverse sedimentation within minutes. This ability is made possible by a reticulated cytoplasmic architecture in *Pyrocystis noctiluca* that enables this rapid increase in overall cell volume without dilution of its cytoplasmic content. We further present a generalized mathematical framework that unifies cell cycle driven density regulation, stratified ecology, and associated cell behavior in the open ocean. Our study unveils an ingenious strategy employed by non-motile plankton to evade the gravitational sedimentation trap, highlighting how precise control of cell size was essential for survival in the ocean.

## Introduction

Plankton, *the wanderers*, owe their namesake to an apparent dependence on the motion of the water surrounding them. Nevertheless, these organisms evolved for billions of years to actively navigate a vertically stratified ecosystem either as large colonies [1] or single cells [2, 3]. These migrators encounter two boundary conditions: one of the sunlit ocean surface, and the other of the darkness at the ocean floor[4–6]. Between these two limits, gradients of temperature[7, 8], pressure[9, 10], salinity, nutrients, and elemental composition[11]act as drivers for biochemical adaptation in planktonic cells.

For photosynthetic organisms, the dilemma of vertical positioning in the ocean arises from the need to maintain proximity to sunlight in the euphotic zone while also gaining access to nutrients that might be available only in deeper waters [12, 13]. Although cell density is tightly regulated amongst most cells compared to cell mass [14], a planktonic life style imposes new constraints on this critical physiological parameter. Given that the eukaryotic cytoplasmic density is typically 5-10% denser than seawater, there is a constant risk of sinking below the euphotic zone and losing the ability to photosynthesize [15]. The evolutionary mechanisms shaping the metabolic and reproductive processes of drifting cells must therefore consider the density of biochemical products and their influence on sedimentation. We term the limit of sinking below the habitable zone for a given phytoplankton species due to sedimentation as a “gravitational trap”. At the very least, a cell must compensate for passive sinking by actively rising to ensure that its progeny do not progressively descend in depth and ultimately sediment to the ocean floor.

Although ciliary and flagellated motility is common, many abundant single-celled phytoplankton, such as diatoms *Coscinodiscus bouvet, Rhizosolenia castracanei*, and *Ethmodiscus rex*, as well as the dinoflagellates *Pyrocystis noctiluca, P. lunella*, and *P. fusiformis*, regularly navigate distances spanning 10 to 100’s of meters without typical motile organelles like cilia or flagella [16–19]. These journeys can be daily, or as in Pyrocystis, ‘once in a lifetime’[20]. This raises the question: without flagella, how do these single cells migrate across length scales hundreds of thousands of times their size?

To illustrate this problem, consider a spherical cell of radius *r* and intracellular fluid of density *ρ*_*cell*_ suspended in ambient seawater with density *ρ*_*sw*_ and dynamic viscosity *µ*(Fig. 1F). In a regime where viscous forces dominate over inertial forces for a suspended cell, sedimentation dynamics is governed by the linearized Navier-Stokes equation [21] giving rise to a simple relationship between cell size, cell density, and sedimentation velocity

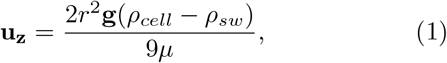

where the gravity direction **g** is aligned with the z-axis and homogeneity and isotropy are assumed in the horizontal *x*−*y* plane. This simple relationship provides a link between cellular physiology(cell size *r*, cell density *ρ*_*cell*_), ecology (defined by the density of seawater as a function of depth *ρ*_*sw*_(*z*)) and cell behavior (either positive or negative sedimentation velocity). Understanding each parameter’s change over time (and depth) would provide a mechanistic explanation linking cell-cycle to ecologically relevant behavior.

**FIG. 1.**
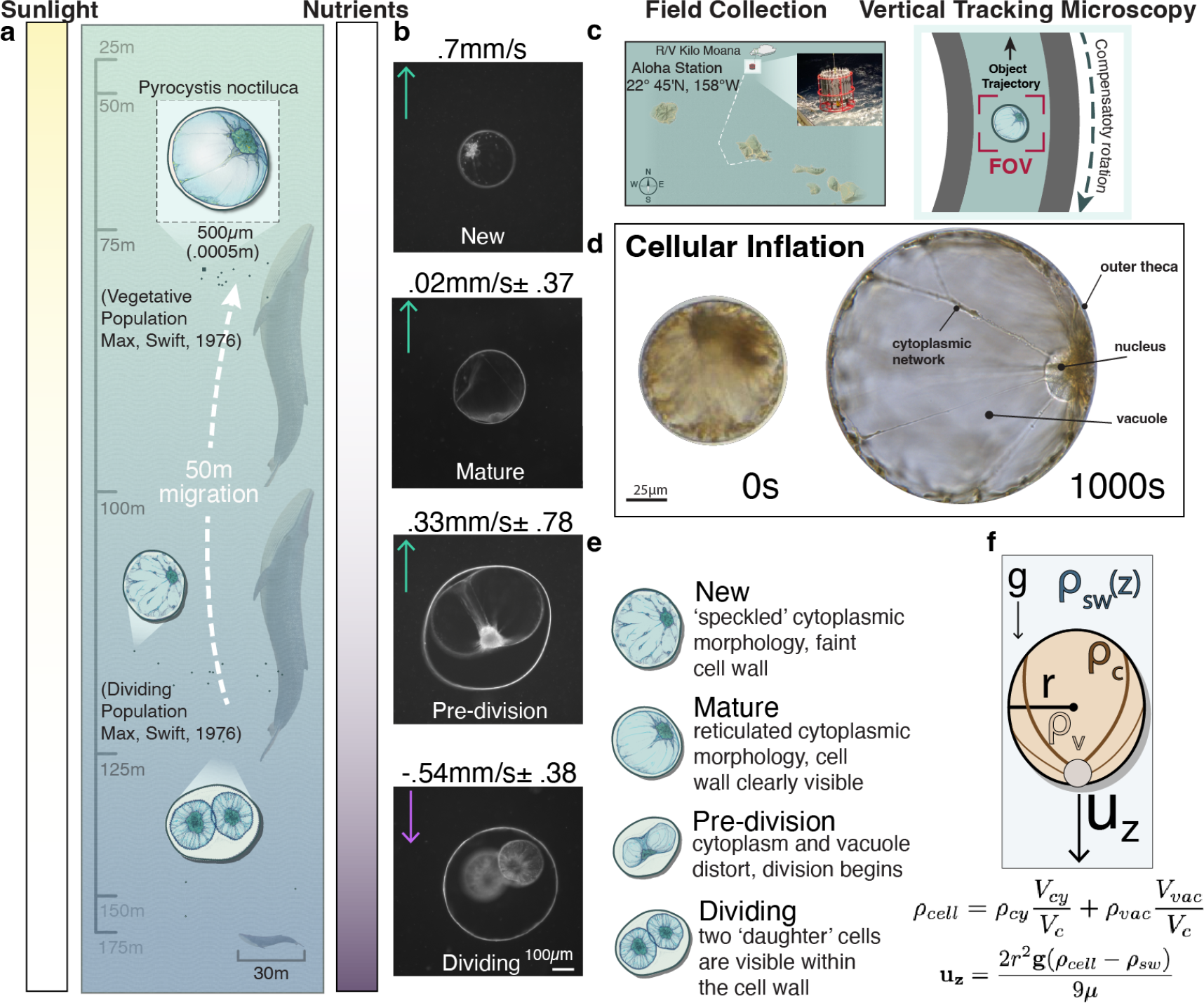
A vertically migrating single cell. *Pyrocystis noctiluca* was collected during the Hawaii Ocean Time series (HOT317) cruise, encountered in deep waters, enabling us to make sedimentation velocity (behavioral) tracks of cells captured and imaged under field conditions. a) *Pyrocystis noctiluca* undergoes a vertical migration of over 50m within the lower euphotic zone. Dividing cell populations are centered 55m below mature cells. Whale bar refers to average length (30m) of a Blue Whale(*Balaenoptera musculus*). b) Wild Pyrocystis cells displayed differing vertical velocities, often associated with morphological differences. c) Cells were collected 100km off the Hawaiian Archipelago on board the RV Kilo Moana using either a CTD rosette or net and loaded directly into the vertical tracking microscope ([29]), where micrographs are collected along with cell trajectories. d) A *Pyrocystis noctiluca* is capable of increasing its volume 6 fold in only 10 minutes, immedietly following cell division. e) Pyrocystis cells can be binned according to cell state, in which both the size and geometry of the inner membrane largly change from (1)newly formed cells,(2) mature cells, (3)cells beginning to undergo mitosis, and (4) cells with two daughter cells present.f)The sedimentation of a spherical cell is a function of its size and density relative to the surrounding medium.

Since the effective cell density (*ρ*_*cell*_) is usually 5-10% higher than that of seawater (*ρ*_*sw*_), this suggests that a cell must actively regulate its effective density to achieve upward motion at some point during its life cycle[15, 22, 23], or else risk sinking to the bottom of the water column. Since sedimentation velocity is a product of cell density and square of cell radius (1), in principle, a cell could internally change either or both its radius (*r*) and density (*ρ*_*cell*_) as a function of the cell cycle, thereby regulating its position in the water column. Important though, is the fact that a closed life-cycle dictates a recurrent occurrence of specific cell size (radius) and cell composition/density at particular time points during the cell cycle (periodic boundary conditions).

Previous studies have explored the potential role of unusually large vacuoles in facilitating buoyancy [17, 24– 26]. Specifically, for *Pyrocystis noctiluca*, it is believed that the cell alters its buoyant mass through regulating the molecular composition of its vacuole. This includes maintaining an abnormally low pH of 3.5 and sequestering compounds which carry a large specific volume — a strategy also employed by a large family of teuthoid squid [27, 28]. However, during replication, cellular mechanisms are reassigned, leading to disruptions in cellular volume, cytoskeletal networks, and transcriptional programs. Unlike some differentiated cells, single-celled organisms must undergo this dramatic rearrangement to reproduce. For plankton, whose size and shape are intimately tied to controlling their position in the water column, this can pose an existential challenge. Despite our current understanding of plant and metazoan cells, the mechanisms determining and maintaining ideal size and shape for single cells, including plankton, remain largely unknown.

In this article, we present the discovery of a spontaneous singular inflation of a cell, made possible by an intricate reticulated cytoplasmic geometry, which endows a cell with an energetically efficient and orientationally robust vertical migration without a canonical swimming gait. In doing so, we establish a theoretical framework based on Stokesian sedimentation, where gravity imposes a selection pressure on non-motile plankton, in the form of mathematical constraints on the cytoplasmic and vacuole densities throughout its cell cycle.

## Results

### Field Measurements

With an aim to unravel the mechanism by which non-swimming plankton in the oceans traverses meters per day [16–19], we began recording the environmental parameters and native behavior of cells within minutes of them undergoing active migration. To capture single-cell motility behavior of the dinoflagellate *Pyrocystis noctiluca* vertically migrating in the ocean we collected cells 100km off the Hawaiian Island archipelago and recorded trajectories in a custom built scale-free vertical tracking microscope [29] on board the research vessel *Kilo Moana* during the Hawaiian Ocean Time-series cruise *HOT317* (Fig. 1A,C). The cells were captured at known depth using CTD (Conductivity, Temperature, and Depth) triggered rosette bottles and quickly transferred to the annular stage filled with seawater collected at corresponding depth. In this manner parameters such as salinity, pressure, and temperature were collected with the organisms at each depth. To quantify motility vectors and velocities of Pyrocystis we used this seawater-filled wheel as a stage, where upward or downward movements of the object are compensated by stage rotation in the opposite direction using a closed-loop tracking system. This constitutes a “hydrodynamic treadmill” for microscale objects(Fig. 1C) [29]. A ‘virtual-depth’ for the object can be assigned from net angular displacement and radial position while sub-cellular morphology is recorded using microscope optics.

We recovered Pyrocystis cells from a CTD cast at a depth of 75 meters, confirming earlier reports of their residency in the lower euphotic zone [24]. We also noted from CTD data that Pyrocystis abundance was centered around the measured pycnocline—where ocean density increases several percent between 75 and 200m, as had been reported previously[24]. Unlike diatoms, radiozoa, and other cells collected at these depths, many mature (vegetative) *Pyrocystis noctiluca* cells were observed to rise. Confirming earlier reports, dividing cells collected at 75m displayed negative buoyancy (sinking) [24](Fig. 1B, Movie S1). Positively buoyant cells moved upward at rates nearing 0.4mm/s, corresponding to a vertical displacement of tens of meters per day. This raises a puzzle: how do these cells, which presumably lack a self-propulsion mechanism, coordinate directional longdistance vertical migrations over their cell cycle?

Since the vertical tracking microscope provides both sedimentation velocity and cellular state through tracking microscopy, we investigated a possible correlation between the two. Indeed, in cells captured at depth and transferred to the microscope, we found that dividing cells consistently displayed negative buoyancy, as previously reported[24, 30](Fig. 1B). Cells with intermediate morphology exhibited average velocities that were positive, in our limited sampling. We moved to the laboratory to construct a more comprehensive map linking cell state to physiological parameters. For instance, cells might maintain different effective cell densities during various parts of their metabolic cycle and use that to strategically position themselves in the vertically stratified fluid mechanical ecosystem.

### Cell inflation and density measurements

In the context of cellular sedimentation, it is useful to consider the cell morphology in context of two separate components cytoplasm and vacuole volume, as well as their respective densities. The effective cell density can be written as

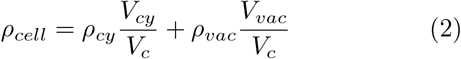

where *ρ*_*cy*_, *ρ*_*vac*_, *ρ*_*cell*_ are cytoplasmic, vacuole and effective cell density, and *V*_*cy*_, *V*_*vac*_, *V*_*c*_ are the volume of cytoplasm, vacuole, and the cell respectively while all of the above parameters might be a function of where a cell is in its cell cycle.

To initiate our study and create a comprehensive physical map, we first focused on visualizing the morphological changes in cultured Pyrocystis cells throughout an entire replicative cycle. On average, cells divide every 7 days in a temperature and light-controlled environment (Movie S3). Long-term time-lapse imaging reveals an extremely dynamic cytoplasmic network, moving material to and from the cell periphery during each day/night transition while undergoing substantial alterations in size and geometry throughout the life cycle (Movie S3). The most significant morphological change in the life of a *Pyrocystis noctiluca* cell occurs immediately following cell division. Each daughter cell is ejected from the mother, often tens to hundreds of microns, then rapidly inflates, increasing its volume more than five-fold in just 10 minutes (Fig. 2A). Even cells forming a presumed gamete instead of two daughter cells, rapidly inflate[31]. Inflation swiftly alters the effective cell size, denoted by cell radius *r*, and can impact effective cellular density *ρ*_*cell*_, depending on the contents of the newly added volume. However, such rapid cellular swelling could have catastrophic consequences on canonical cell physiology, diluting biochemical interactions and considerably straining cell functions [32]. How do *Pyrocystis noctiluca* cells suc-cessfully incorporate this event, and what are the associated consequences for their behavior? The usual process of cell growth is carefully regulated to align with general metabolic timelines, enabling efficient protein production and energy storage [33]. This suggests that any anomalous, inflation-driven growth in this dinoflagellate must have evolved to confer some advantage in its life history.

**FIG. 2.**
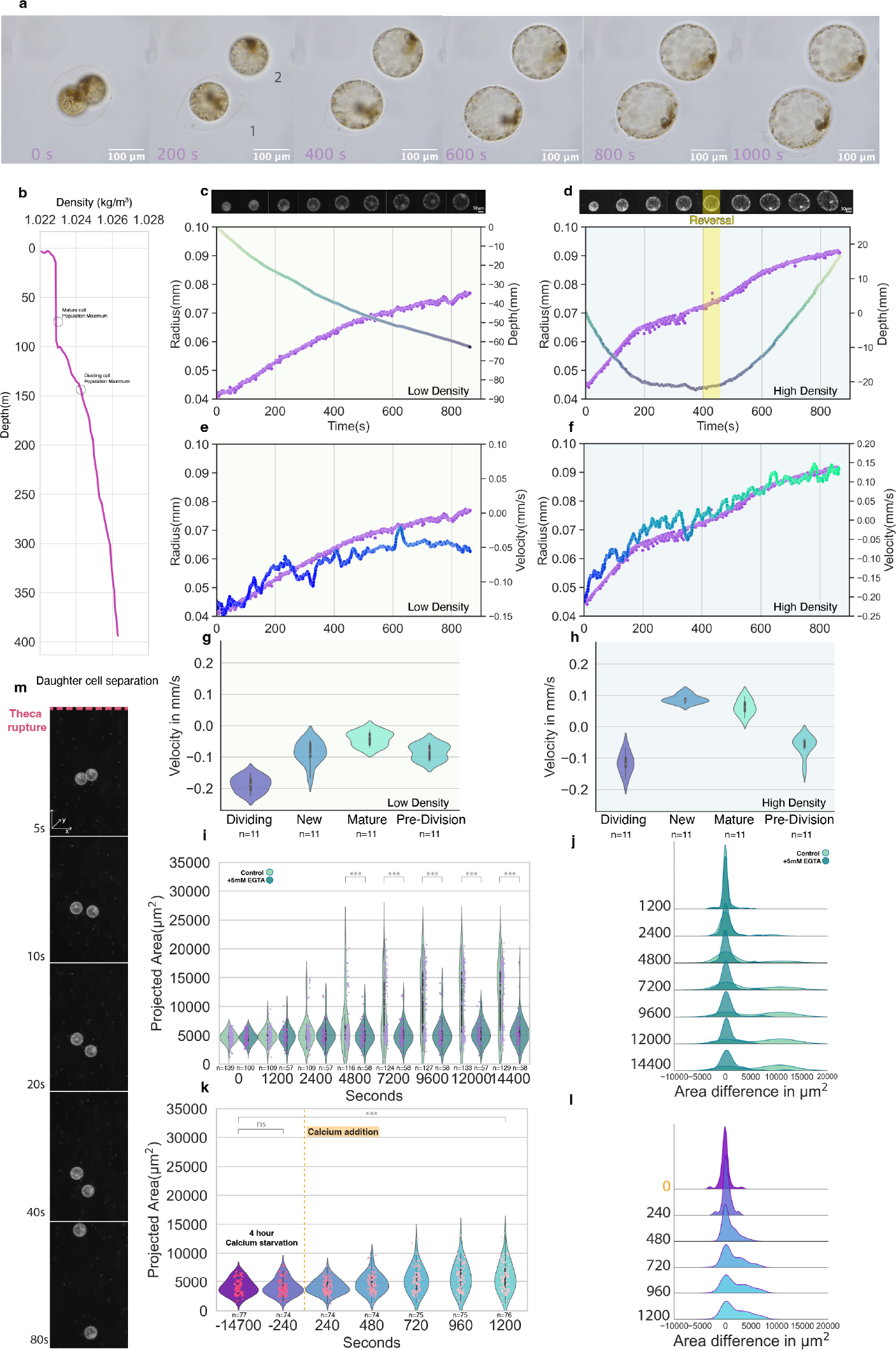
Triggered cellular inflation as a means of motility. a)Post-division, daughter cells undergo an expansion event increasing their volume over 5 fold in 15 minutes.b)Pyrocystis cells migrate along the lower euphotic zone, where dividing cells experience seawater at a density much higher than vegetative counterparts(CTD measurement from Station Aloha 12.19.2019). c,e)Radius increase rapidly decreases sedimentation velocity, with cells switching from sinking to rising at the higher seawater density. d,f)Vertical tracks from individual division events in the gravity machine show a concomitant decrease in sedimentation velocity as the cell radius increases through inflation, with a complete reversal of trajectory in seawater of higher density. g,h)Pyrocystis cells at different cell cycle stages display differing average velocities. i,j) Areas of individual segmented and tracked daughter cells during the 4 hours post dark/light transition. The inflation event is inhibited by addition of the calcium chelator EGTA. k,l)Calcium starved cells are still viable, as cells grown in artificial seawater without calcium undergo expansion on addition of 2mM calcium to the media. m)Microscopy frames following division show daughter cells sediment at different rates, due to residual drag from the mother cell theca on one cell. Time is seconds since theca rupture. The width of the violin plots indicate the density of data points at different values of projected area.The white dot represents the median (50th percentile), the thick black line indicates the interquartile range (IQR; 25th-75th percentiles), and the thinner black lines show the lower and upper adjacent values.*** represent p-value of *<*.001 using student’s t-test.

Our field observations showed that native populations of Pyrocystis were predominantly located near the pycnocline. This region in the water column is characterized by a significant change in seawater density over distances of tens of meters. This observation prompted us to investigate the relationship between cell state (categorized by cytoplasmic/vacuole geometry) and cell density in Pyrocystis cells. Under our laboratory conditions using supplemented artificial seawater(F/2-si), Pyrocystis cultures are viable across a range of salinities. However, under our temperature and light conditions they displayed optimal replicative growth at a salinity of 36 ppt. This salinity also made long-term microscopic observation possible, as in higher salinities/densities many cells are positively buoyant and float off the coverslip(Movie S11). As rapid salinity changes can disrupt cell morphology, to simulate sedimentation through increasing densities on the hours-to-days timescale we used small amounts of Optiprep, an isotonic, non-toxic medium, to maintain cultures in isotonic seawater with varying densities representing different depths in the water column (Fig 2G,H)[17]. Cells maintained in Optiprep supplemented medium showed no defects in growth or morphology, even over several weeks to months. We measured average sedimentation velocities using the vertical tracking microscope and grouped them according to cell-cycle state. Consistent with our previous observations, sedimentation values were most negative in dividing cells. New and vegetative cells were more buoyant, particularly evident in the high-density seawater. A large density change appears to occur within minutes between cell division and the emergence of the ‘New’ cell. To directly observe this process, we recorded cell division events using the vertical tracking microscope (Fig. 2C,D,E,F). In seawater with a density of 1.028 kg/m^3^, cells containing two daughter cells had sedimentation rates of around -0.2 mm/s. After the outer thecal breakdown, the two daughter cells separated and began to inflate. One cell rapidly falls away from the mother cell, with the other slowed down by the ‘hydrodynamic parachute’ effect of the mother cell theca (Fig. 2M). As each daughter cell sediments, inflation coincides with a rapid decrease in cell sedimentation velocity from 0.2 mm/s to 0.1 mm/s over the 10-minute volume increase. Simulating the lower edge of the pycnocline, in seawater with a density of 1.032 kg/m^3^, this event causes the cell to rapidly switch from sinking to rising in the water column (Video S12, Fig.2C). Pyrocystis cells actively dividing in the ocean tend to be located at depths of 50-75 meters below the density maximum of vegetative cells at the denser edge of the pycnocline and near the bottom of the euphotic zone [24].

This density difference can also be observed in bulk, using a density based sorting approach[34]. When mixed cultures of Pyrocystis are passed through such a sorter, the dense fraction is composed of 80% dividing or predivision cells. Most mature cells sort into the less-dense fraction(Fig. S6).

According to Equation 1, sedimentation velocity is directly proportional to the product of the square of cell radius and effective cell density. The decrease in sedimentation velocity as cell radius increases implies that effective cellular density must decrease dramatically. A cell needing to rapidly reduce sinking velocity could employ such a strategy to avoid prolonged residence in lowlight conditions after free-fall during cell division, when any bladder lift is significantly reduced.

### Cellular topology

Tracking demonstrates a direct influence of cellular inflation on Pyrocystis movement, but the physiological mechanisms enabling such a feat are not immediately apparent. To understand how cell physiology might facilitate rapid cell inflation, we visualized individual Pyrocystis cells using electron and light microscopy. Pyrocystis noctiluca has long been observed under light microscopy to exhibit a distinctive cellular architecture characterized by cable-like structures that radiate in a network pattern throughout its expansive and sparsely populated cell body. Yet, the significance of the structure remains unknown. Upon examining individual Pyrocystis cells with a transmission electron microscope, we observed a large, empty vacuole surrounding a membrane-enclosed filamentous network. This network appeared to contain a highly dense mixture of various organelles (Fig. 3D, S3). The entire cellular machinery seems to be constrained to a minimal volume determined by the fiber structure (Fig. 3E inset). We confirmed that the filamentous network observed under the transmission electron microscope contains a highly autofluorescent mixture filled with plastids, as evidenced by wide-field fluorescence images using red and far-red excitation (Fig. S1E, G). Furthermore, we find this cytoplasmic network is not only present on the vacuole surface but also penetrates it at multiple locations rejoining back with the nucleus and linking these two organelles. To quantify the volume fraction of cytoplasm and vacuole, we employed volumetric fluorescent microscopy. We found that the reticulated cytoplasmic network connecting the nucleus with the periphery constitutes less than 5% of the total cell volume (Fig. 3A, S3A). However, pre-division daughter cells are much denser, with the cytoplasmic network making up a significantly larger portion of the total cell volume (Fig. 3B, S3A).

**FIG. 3.**
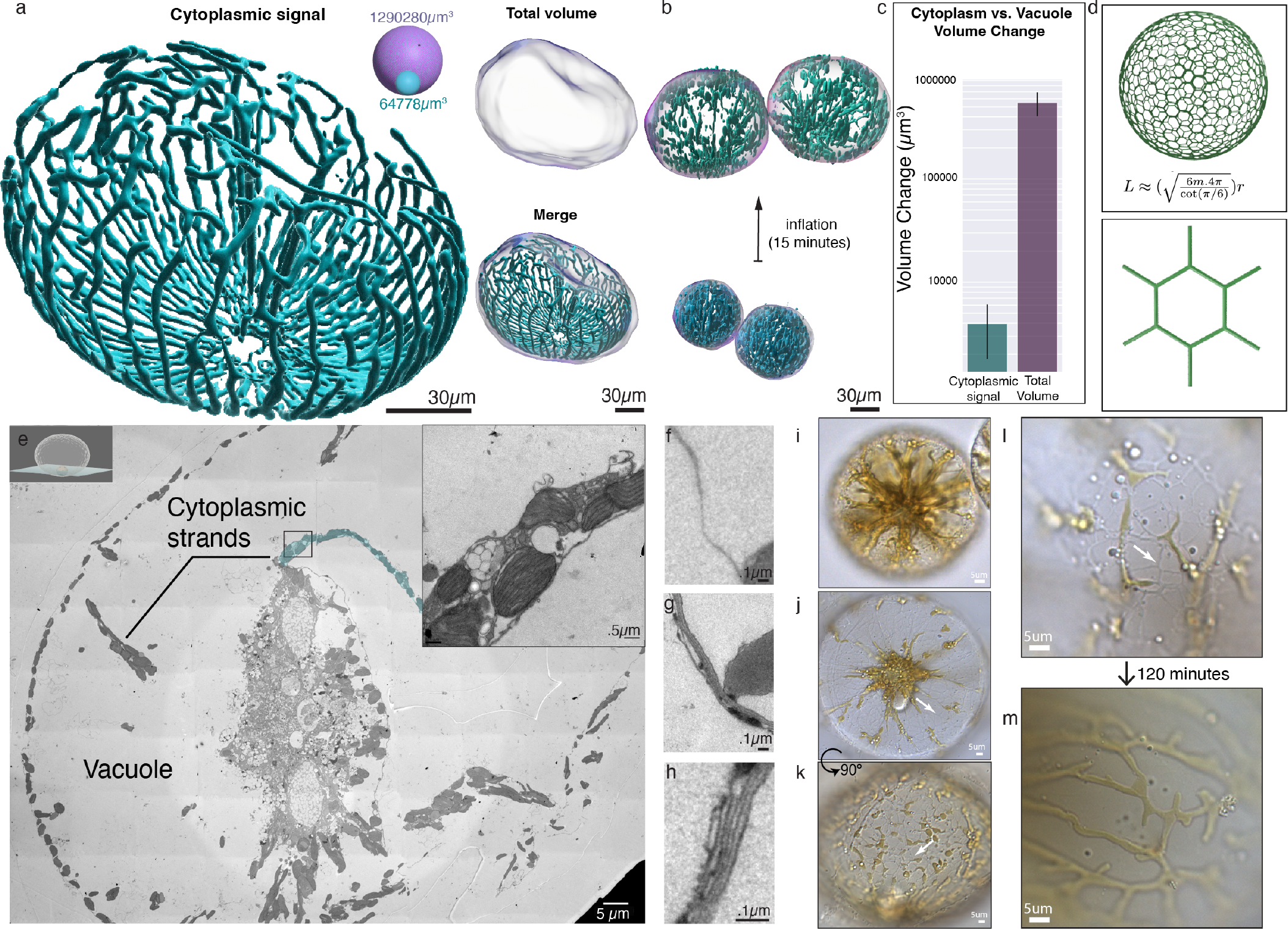
Cellular architecture in 3D. a)Pyrocystis’s cytoplasm is organized into a reticulated network that surrounds, and transects, the large inner vacuole. Inset visualizes relative volume difference between total cell volume and the cytoplasm, which makes up 5% of total cell volume b,c) Quantification of cytoplasmic and cellular volume change before and after inflation in 3 pairs of dividing cells. ‘Cytoplasmic signal’ refers to cytoplasmic autofluorescence signal, ‘Total Volume’ refers to cell volume encompassed by the cell network. Error bars represent 95% confidence interval. d) This cytoplasmic structure can be described as an ‘n-torus’, where ‘n’ is the number of holes in the manifold-in this case the membrane bound cytoplasm. Such a structure allows for large cell volume changes without dilution or concentration effects on cytoplasmic contents. e) Electron micrographs display the highly jammed composition of the cytoplasmic network. Inset highlights the membrane-bound organelles within a cytoplasmic filament. f) A mature cell vacuole is bound by a single contiguous membrane. (g,h) The membrane in an uninflated daughter cell appears as a laminated structure of 5-8 membrane layers. This could enable large and rapid increases in surface area. i)Pre-inflated daughter cells are more compact, with cytoplasmic contents compressed into several ‘pillars’.j,k,l)In recently inflated cells, the membrane network is clearly seen with and without pigmented cytoplasm inside.m)As the cell matures, most of the membrane network is filled with cytoplasm.

Through live cell imaging of the inflation process, we found that Pyrocystis cells undergo a dramatic increase in total volume, while the cytoplasmic signal remains nearly constant (Fig. 3B,C, S3A, Movie S6). These observations suggest that the cell inflation process involves an increase in vacuole size, rather than an increase in cytoplasmic volume.

Pyrocystis cells present a unique cellular architecture and topology, an “n-genus torus” linked to a spherical vacuole. We examine the implications of this topology to cellular inflation. As described previously, a six-fold increase in cell volume in less than 10 minutes puts an immense strain on cellular geometry. As cell radius increases rapidly due to inflation, the overall vacuole surface area would need to increase proportional to the square of its radius. But a reticulated cytoplasmic network architecture, linked to this vacuole, due to it’s unique topology, would only increase linearly in its length (see SI for details). This significantly reduces necessary material. Furthermore, this inflation can be achieved without any dilution of the cytoplasm of the cell. This topologically complex cellular architecture enables a fast cell inflation event without putting a significant constraint on the required cell-membrane that would otherwise needed to be generated in a short period of time.

Topologically, the division of an n-torus cell also presents several challenges because of this unique membrane architecture. Importantly, these rearrangements take place within the confinement of the “outer theca” wall which largely conserves total apparent cellular volume and hydrodynamic profile while the associated vacuole and entangled cytoplasmic network divide(Fig. S1A) [35]. The topology undergoes a complete rearrangement at the point of cell division. After partitioning into two distinctly visible cells inside the theca, we observe a rapid breakdown of the wall. This dissolving wall eventually bursts and ejects both daughter cells(Movie S4,S5). This separation is immediately followed by a rapid inflation of both daughter cells, which increase their volume 3-6 fold over 15 minutes(Fig. 3b). The somewhat ‘speckled’ nature of the fluorescence signal is visualized well by high magnification light images of the reticulated membrane network(Fig. 3I-M). Inflated cells display a cytoplasm that is randomly distributed throughout the network, with ‘empty’ membrane tubules clearly evident(Fig. 3I,K). As the cell matures, these tubules become more completely filled, forming an interconnected network, with few ‘unfilled’ capillary bridges.

For the vacuole and the cytoplasm to persist together despite inflation and division cycles, these two topological objects must remain linked together. This is clearly visible as cytoplasmic tunnels that pass through the vacuole geometry from one side to another. This can be further quantified as a linking number that ensures that no deformation of the geometry of cytoplasm or vacuole can separate the two.

### Extracellular calcium triggers inflation

How is such a dramatic change in cellular architecture orchestrated? For cells, volume regulation is a critical aspect of cellular homeostasis, controlled by various families of pores, channels, and pumps within different signaling networks [36, 37]. We used known channel interfering agents to investigate the mechanisms involved in inflation.

We recorded inflation in cell populations at the beginning of their conditioned light cycle and quantified it using projected 2D cell areas as a proxy for volume. In seawater with a salinity of 36 ppt and specific gravity of 1.028, inflation resulted in approximately 2.5-fold growth in projected area over a 240-minute observation period. Supplementing the artificial seawater medium with the calcium chelator EGTA or channel blockers like Cobalt completely inhibited inflation (Fig. 2I,J, Movies S7,S8) (Fig. S2). Cells in isotonic seawater prepared without calcium also did not inflate (Fig. 2K). Notably, these cells remained viable; upon adding a bolus of calcium chloride solution to the medium, inflation commenced within 60 seconds and completed within the typical 15minute window (Fig. 2L). The final inflated size was smaller than that of calcium-replete cells, which is expected given calcium’s role in other crucial cell activities. These cells also remained competent for further rounds of cell division when allowed to continue their life cycle in the presence of calcium. The absence of calcium ions in replete seawater or in calcium-free artificial seawater rendered the inflation mechanism inactive, suggesting that the process is actively regulated by components in the cell membrane.

Theca breakdown, which takes about an hour, is also timed to inflation. If the cell wall is compromised with a micro-pipette before natural breakdown, exposing the daughter cells to seawater, inflation begins prematurely. These daughter cells conform to the mother’s theca profile until enough internal pressure is built up to escape (Movie S10). These two processes are linked to ensure a timely reversal of sedimentation.

Diffusive processes are often insufficient for the translocation of large volumes of water across membranes, yet many biological processes require such flux [38, 39]. In metazoans, processes involving ionic flux on a large scale, such as renal function, rely on channel proteins called aquaporins, which can transport up to 3×10^9^ molecules of de-ionized water per second [40, 41]. Expression of aquaporin protein in a Xenopus oocyte can cause these millimeter-sized cells to swell over 1.5-fold within minutes before rupturing [42]. It has also been observed that aquaporins’ spatial expression in cancer cells creates an ‘osmotic engine,’ allowing for directional movement through tight junctions [43].

Plant processes, such as pollen tube elongation and leaf movement, also require selective movement of deionized water [44–47]. Pollen tube elongation, another example of rapid cell expansion, is controlled by a calcium gradient. Given aquaporins’ potential to facilitate the coordinated influx of low-density ballast, we sought to confirm their presence in dinoflagellates.

For this we assembled a working de-novo transcriptome for *Pyrocystis noctiluca* using Iso-seq long-read sequencing. When compared to known aquaporins across the tree of life the *Pyrocystis noctiluca* transcriptome encoded several proteins identified as aquaporins(Table. 1, Fig. S4). Using AlphaFold for protein structure prediction, several homologous sequences revealed canonical 6pass Aquaporin structures (AQP1) along with other conformations such as the pollen-specific NIP4 [48]. NIP4like hits are intriguing as these proteins are crucial to the hydration and vacuolation of pollen during development, a process requiring rapid volume increase [47](Fig. S4). The presence of these channels and numerous mechanosensitive, voltage-sensing, and rectifying channels display a capable toolkit for size regulation in dinoflagelletes.

This lays the foundation for unraveling molecular players and regulatory processes that enable timely cellular inflation.

### Cells evading the gravitational trap

Our ecological and physiological measurements allow us to construct a theoretical framework to understand how a single cell navigates ∼10^5^ times its own size without swimming. Unlike swimming at low Reynolds numbers [49, 50] where changes in the configuration of an organism leads to swimming strokes, in the non-motile cell under consideration activity manifests through changes in its internal state (cell cycle clock) which leads to a dynamic buoyancy. We construct an equation of motion for the trajectory of a single cell, with a dynamic buoyancy during the cell cycle; and find that rapid inflation contributes a singular perturbation to the vertical migration dynamics.

The key aspects of this derivation involves – mapping time coordinate to cell cycle phase (*t*→*ϕ*), writing an analytical expression for effective cell density *ρ*_*cell*_(*ϕ*) and cell radius *r*(*ϕ*) over the entire cell cycle using the mathematical technique of asymptotic matching [51] which connects widely separated length and time scales, and substituting this density in the Stokes law (1) to construct a differential equation (4) that describes the trajectory of a single cell in a vertically stratified ocean over multiple cell cycles, where oceanic density stratification is captured by allowing seawater density *ρ*_*sw*_ to be a function of ocean depth *z* . As will become transparent in the calculation below, we non-dimensionalize distance and time using the characteristic sedimentation length and time scale in the system. We now begin presenting these steps elaborately.

From (1), cellular behavior (speed of rising and sinking) is a function of cell size, cell density, and surrounding sea water density. These parameters change during a cell cycle – radius and density changes due to internal changes in physiology of the cell as the cell-cycle clock proceeds, and the ambient seawater density changes due to the stratification of the ocean. We define a non-dimensional cell phase variable *ϕ* = *αt/nT*_*c*_ where *t/T*_*c*_ is the time scaled by the total time period of the cell cycle (∼7 days) and *n* = 1, 2, 3.. is the number of cell cycles. We have introduced activity through a constant parameter *α*, which drives the cell cycle at a constant rate *α/T*_*c*_. Thus all the internal degrees of freedom of a cell are dependent on *ϕ* which quantifies the stage (phase) of a cell is during its cell cycle. We depict the relevant phases of the cell cycle(Fig. 4A), by arbitrarily assigning *ϕ* = 0 at the start of cell inflation (starting at the moment two daughter cells break open out of the thecae of mother cell) and continuous through cell growth, pre-division, and duplication. *ϕ* is reset as a cell completes its cell cycle in *T*_*c*_. Since *α* can be absorbed in rescaling *ϕ*, henceforth we set *α* = 1 throughout this analysis, while endorsing the presence of an activity parameter in this system. Note that in general the cell cycle phase *ϕ* could be a non-linear function of time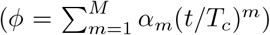, with multiple activity parameters *α*_*m*_ driving the cell cycle; however, for the sake of simplicity we do not discuss this rich plausibility further and assume that the cell cycle clock ticks at a constant rate.

**FIG. 4.**
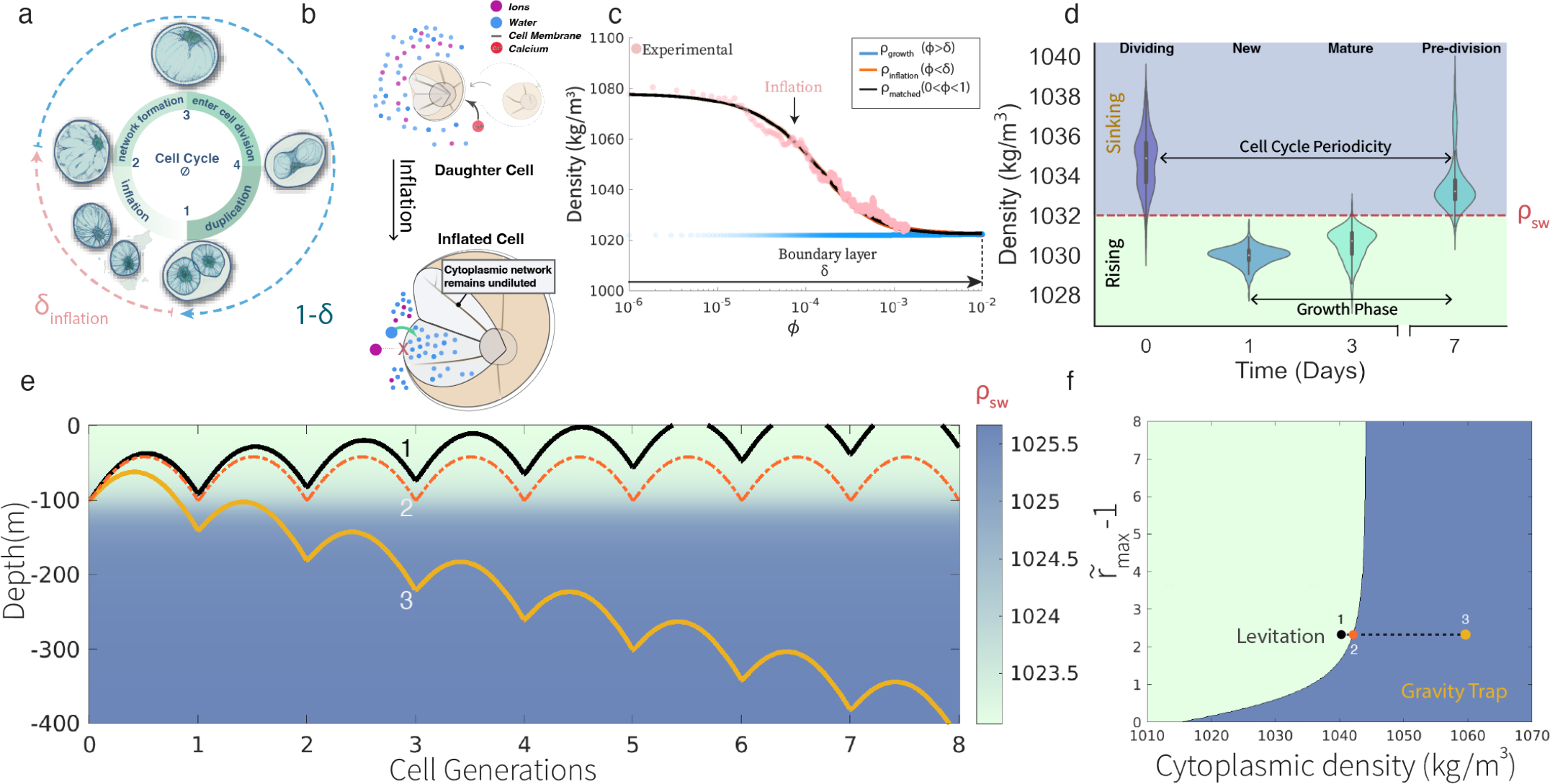
A minimal model for inflation-based self propulsion. a)Pyrocystis undergoes numerous geometrical membrane transformations throughout its life-cycle while maintaining a general ellipsoidal form-factor to its cell-wall. 1.On cell-wall breakdown daughter cells quickly expand. 2. This material becomes organized into a reticulated network over the next few hours. 3. The mature Pyrocystis cell maintains a large vacuole and distributed cytoplasmic network for around 96 hours. 4.Division begins with a reorganization of the network and vacuole. This ‘bowtie’ shape eventually forms 2 spherical daughter cells. b)Proposed role of selective molecular import in escaping a gravity trap. At the completion of cell division, the two daughter cells have greatly reduced vacuoles compared to mature cells. Without positively buoyant vacuole material they sediment to the lower euphotic zone. An unknown cue induces cell wall breakdown which then exposes the membrane to seawater. Calcium then binds and activates channels, including aquaporins, to import water into the vacuole. This rapidly increases cell volume while greatly reducing density and reversing sedimentation velocity. c) Comparing the asymptotically matched effective density in Supplementary equation 10, with the experimentally observed density in the inflation phase, gives *ρ*_*cy*_ = 1078*kg/m*^3^ and *ρ*_*vac*_ = 1015*kg/m*^3^ with a boundary layer of thickness *δ* = 4 *×*10^*−*4^ and 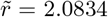. d)Experimentally measured density values over the cell cycle. e)The trajectory of cells for various cytoplasm densities. The yellow trajectory is that of a cell with cytoplasmic density beyond the critical density, wherein inflation cannot overcome the gravitational trap, thus it bounces down into the abyss in multiple cell cycle times, the black trajectory represents a cell that rises in a water column with successive cell division, and the red trajectory rests at the critical density, generationally bouncing between sinking and rising within a range of ocean depth. f) Phase diagram with a critical density boundary beyond which even extreme inflation cannot counter downward gravitational pull, defining the gravity trap (blue region). The colored circles correspond to the levitating, rising, and trapped trajectories in D) respectively.

We map the time in equation (1) to the phase coordinate (*t*→*ϕ*) and non-dimensionalize (1) using the length of the daughter cell *L* = *r*_*min*_, and the time scale *T* = 9*µ/*(2*g*(*ρ*_*cy*_−*ρ*_*vac*_)*r*_*min*_). Redefining 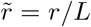 and 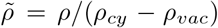 gives the equation of motion in 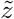 and *ϕ* plane (defined as the solution space for where a cell is at depth *z* as a function of its cell cycle phase *ϕ*)

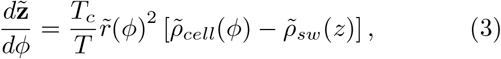

where the effective density 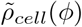 given by the nondimensionalized expression (2), is a function of the internal state *ϕ* of the cell (Fig. 4A) and is independent of the precise morphology of the cytoplasm; which indeed has a non-trivial architecture (Fig. 3), and *ρ*_*sw*_(*z*) captures the pycnocline – the increase in sea water density as a function of depth *z*. Thus (3) can be seen as describing the dynamics of cell behavior captured in cell height, as a function of a complete cell cycle in an ocean sculpted by the pycnocline. Although fluid turbulence is universally present in marine ecosystems [52], turbulent mixing in the vertical direction is suppressed in regions away from the boundaries of the fluid (air-water and land-water interface) [53]. With the current focus on understanding the deterministic part of the dynamics [equation (3)], we assume that the vertical flow velocities are negligible.

In our experiments we measure settling velocity and size, and get the effective cell density *ρ*_*cell*_ using (1). We construct an analytical expression for *ρ*_*cell*_(*ϕ*) in equation (3), that captures the experimentally measured densities, by considering two critical stages of cell cycle where large scale density change occur (Fig. 4 A) – first, the fast time scale (*T*_*B*_*/T*_*c*_) of rapid ballooning during which the cell density drops sharply due to selective intake of water in the vacuole (Fig. 4 B, C), and second, the slow time scale (1 −*T*_*B*_*/T*_*c*_) process of cell growth where cell mass accumulates via production of cytoplasmic content, making the cell more dense [see Figure 4 D]. In the second phase, the density grows gradually to the same value that the cell had prior to its division, as observed in our experiments [see Figure 4 D], for daughter cells are expected to be identical to the mother cell in “all” respects; a periodicity that is crucial to the long term dynamics of the cell. We mathematically stitch these two widely separated time scales using asymptotic matching of the density at *ϕ* = *δ*≡*T*_*B*_*/T*_*c*_, which yields an expression for *ρ*_*cell*_(*ϕ*) that is valid throughout the cell cycle [for a detailed derivation see Supplementary text]. The resulting Stokes equation in the 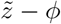 plane takes the form

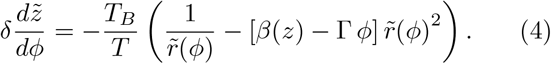

Here *T*_*B*_*/T* is the ratio of ballooning time scale to sedimentation time scale, 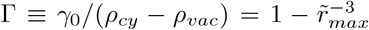 is the measure of relative density reduction due to ballooning. A non-dimensional density parameter *β*(*z*) = (*ρ*_*sw*_(*z*) −*ρ*_*vac*_)*/*(*ρ*_*cy*_ − *ρ*_*vac*_) captures the external environment through the pycnocline – the increase in sea water density *ρ*_*sw*_ as a function of depth *z*, and internal cell parameters through vacuole density, and the cytoplasmic density. We have thus constructed a single differential equation which captures the settling behavior throughout the cell cycle, with simplifying assumptions about the relevant stages of the cell cycle. This minimal 1D model incorporates a wide range of length (cell radius approx. 50 microns to ocean stratification depth approx. 100 meters) and time (ballooning approx. 10 mins to cell cycle approx. 1 week) scales, while coupling the physical and biological components of these ecophysiological dynamics.

In equation (4) a small parameter *δ* multiplies with the derivative (*δ*∼10^−3^ in experiments), and therefore the ballooning event can be mathematically treated as a singular perturbation to the dynamics [51]; thanks to the “abrupt” ballooning of the cell [see Fig. 4 B]. It is thus amenable to a singular perturbation analysis in which the inner (inflation phase) solution, to leading order in *δ*, is the solution to

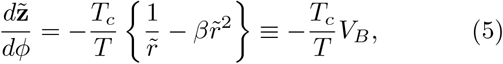

which has a fixed point in non-dimensional velocity *V*_*B*_ = 0 at 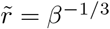, at which cells become neutrally buoyant, implying that via ballooning (increasing 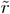) cells can steer themselves to rise from a sinking fate; consistent with our field observations(Fig. 4 A-E). Since the lifetime of the ballooning event *δ* is negligibly small compared to the total transit time of the organism, only the outer (growth phase) solution of equation (4) is ecologically relevant. This singular perturbation in density generates a cusp singularity (via bouncing) in the trajectory of the cell in 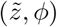 plane [see Figure 4 E], which remains elusive to field observations due to its fast nature. In a constant pycnocline approximation, the outer solution to (4), to leading order in *δ*, becomes

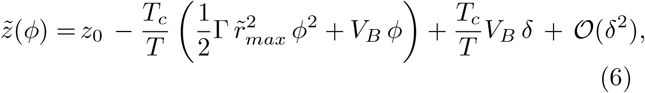

where *V*_*B*_ is the velocity immediately after ballooning, given by (5) with 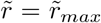[see Fig. 2 E], and *z*_0_ is the ocean depth at which the first cell cycle begins. The ecophysiological and mathematical relevance of ballooning based migration also enables resetting of the “slingshot” and adding residual corrections to the trajectory as a perturbation expansion in *δ*. The notable outcomes of this analysis are:

1. *Criticality* – equation (6) is a parabola in (*ϕ,z*) plane with the vertex at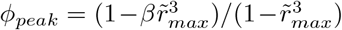). When *ϕ*_*peak*_ = 1*/*2 the cell returns to the same vertical height as it started during division, thus setting *ϕ*_*peak*_ = 1*/*2 gives the critical ballooning radius *r*_*c*_ required for escaping the gravity trap (gradually rise in the long time scale of multiple cell cycles). The critical radius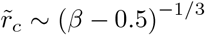, diverges as *β*→ 0.5 with a critical exponent 1*/*3, giving the condition

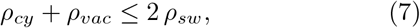

on relevant densities for an escape. This implies that beyond a set of critical parameters it is not possible to escape the “gravity trap” and eventually a cell will sink if no upwelling brings it up (Fig. 4C, D), thus formally allowing gravitational force to apply selection pressure in the evolution of phytoplankton in this framework.
2. *Effective inertia* – for a constant pycnocline, equation (4) can be mapped to the equation of a projectile shot vertically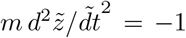, with an effective mass or inertia 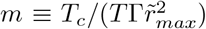) which depends on the ballooning radius and effective drive *α* = 1 which is defined as a constant force that drives the cell cycle clock in Fig. 4A. This effective inertial dynamics in a Stokesian regime comes from internal degrees of freedom actively driven in the cell. This seemingly exclusive possibility also appears in the dynamics of micro-swimmers [54, 55] and passively sinking particles with internal degrees of freedom [56]. To draw the *z*−*ϕ* phase in its completeness we numerically integrate (4), using the pycnocline measured by CTD during the HOT 317 cruise, *ρ*_*sw*_ = 1024.3 + 0.9 tan^−1^ 0.05(*z* −107.8)*kg/m*^3^, and assuming only fresh water permeates through the membrane *ρ*_*vac*_ = 1000*kg/m*^3^. We find escaping and trapped trajectories in Fig. 4C, for small and large values of cytoplasmic density *ρ*_*cy*_ respectively. This clearly demarcates a space of parameters that can lead to a cell being perpetually suspended while navigating the water column vertically as compared to another cell population which will eventually sink to the bottom of the ocean. Note that in our calculations this phase space becomes unbounded and rises sharply around the cytoplasmic density 1040*Kg/m*^3^(Fig. 4 f). This critical behavior does not depend on the initial vertical position of the cell, thus the phase diagram is robust to the vertical flow fluctuations in the ocean. The ballistic motion resulting from the dynamics of buoyancy control presents a unique advantage for organisms that demonstrate long-distance migrations such as Pyrocystis cells. Because this mechanism is robust to any orientational noise/diffusion and drift caused by fluid shear, organisms can maintain course over long distances as compared to classical swimmers that are highly sensitive to cell orientation. The framework presented above can be extended to a number of aquatic organisms experiencing a downward pull and provides insights into organization of cell cycle in context of vertical migration. Thus ballooning in a cell in context of a vertical migration can be considered as a necessary *slingshot* mechanism needed to escape the “gravity trap”.

## Discussion

Our physiological and kinematic measurements allow us to identify a critical piece in the puzzle of a nonflagellated swimmer: cellular inflation. Upon the completion of a cell cycle, the cell initiates water influx through a calcium-activated mechanism, swiftly altering its density relative to the surrounding seawater. This single parameter determines whether the cell moves up or down, and how quickly. Inflation has several features that are well-suited to a long-distance vertical migration. Water movement at the time of inflation is energetically passive-following the concentration gradient. This approach allows the cell to avoid expending energy for growth at its maximum depth, where light(and energy)is scarce. Additionally, it offers a vertical trajectory that is independent of lateral motion or orientation. This process demonstrates how precise control of cell physiology can serve as an alternative to the flagellated locomotion. The present theoretical framework offers a dynamical system perspective to the life of a seemingly non-motile plankton suspended in the ocean; where motility comes from active buoyancy control rather than a conventional swimming mechanism [49, 50]. Our theory stitches together widely separated length and time scales, resulting in a minimal 1D model that incorporates both the physical and cell-physiological aspects of this reduced ecosystem. It offers a framework in which gravitational pull can formally operate as a selection pressure in the evolution of marine plankton [density inequality as in equation (7)]. In future work, the dynamical equation (4) can be further embellished to account for a broader group of ecological parameters including variation in salinity, temperature and an explicit cellular feedback. Although the mathematical forms of *ρ*_*inner*_ and *ρ*_*outer*_ [See Supplementary equation 7 - 10] comes from observations specific to *Pyrocystis noctiluca*, the matched asymptotic approach presented here is likely generalizable to a richer class of plankton with cell cycle phases involving multiple boundary layers, and non-linearities in the growth phase. The current model presents a deterministic dynamical system. Since fluid turbulence is a universal feature present in all marine ecosystems, we can further incorporate it in our models either as stochasticity in equation (4) or more elaborately by coupling equation (4) with the Navier-Stokes equation for the fluid flow [52].

The planktonic migrations that drive the ‘biological pump’ represent the foundation of the most significant daily movement of biomass on Earth. The evolution of precise control over a single cellular parameter, *ρ*_*cell*_, has facilitated the development of a remarkably efficient alternative to traditional mechanical swimming motility. This ingenious adaptation takes advantage of the critical features intrinsic to the oceanic ecosystem. Further exploration of these mechanisms may not only enhance our understanding of cellular and evolutionary processes but also inspire solutions to various challenges in engineering that draw from the natural world.

## Methods

### Obtaining dinoflagellates

Pyrocystis cells were first obtained from a CTD rosette bottle triggered at a depth of 75m. When foul weather prohibited further casts, vertical plankton tows were done from 25-35m with a 20 micron mesh plankton net at a max speed of 1kt. Presence of Pyrocystis noctiluca in the sample was confirmed via imaging on a flow-through Planktoscope instrument. Cells were then isolated on a dissection microscope and placed in filtered underway water. 5 micron polystyrene beads were added to the media prior to loading into the sample stage to mark flow fields for later PIV correction. **Cell Culture** Pyrocystis noctiluca (CCMP732) were ordered from Bigelow Labs and expanded in F/2 media(Si) prepared using artificial seawater (reef salts) at a salinity of 36%, specific gravity 1.028. Cells were cultured at 21ºC under cool white fluorescent light at 2500 lux. Media was refreshed every 2 weeks to maintain proper salinity. Cells were expanded every 6 weeks to minimize accumulation of cell debris. For seawater density increase, a solution of Optiprep/2X seawater was prepared and added as needed to create isotonic seawater of higher density. Density was verified in a refractometer instrument.

### Pyrocystis division assay

Dividing Pyrocystis cells (with 2 daughter cells visible inside the cell wall) were isolated on a dissection scope, washed in fresh media, and placed in 5ml of F/2. Mature cells were included as a proxy of cell viability, as plasmolysis is much more apparent in the mature cells. EGTA was added from a .5M stock solution. DI Water was added as mock to control cells. For the calcium depletion experiment, cells were sorted, washed, and placed in F/2 media prepared with calcium-free artificial seawater prepared according to the Cold Spring Harbor protocol. The calcium bolus was from a .5M CaCl stock solution.

### Image Processing

*Bulk Assay* To quantify the area increase in a population of dividing cells, a custom model was generated within Cellpose v2.0. This was done by updating the pretrained ‘Cyto2’ model with an annotated mask layer drawn in the Napari viewer containing 150 dividing Pyrocystis daughter cells. Cells were then morphologically categorized according to cell state. Dead or Plasmolysed cells were excluded from analysis at this stage. Unique labels were then applied to each daughter cell in the mask using OpenCV. A particle tracking code was written and employed to follow single daughter cells through the time-series and plot their area increase over time. Projected cell area was then calculated using the region properties function in OpenCV. *Vertical Tracks* To quantify the area in sedimenting cells the Watershed algorithm was applied using OpenCV after simple morphological operations to remove background. In frames where the cell drifted out of focus in Y, no volume change was assumed between the current and previous frame. All code, including models, is available in the public repository on GitHub.

### Vertical Velocity Measurements

To quantify the vertical velocities of cells in the tracking microscope, background flows from thermal fluctuations or other movement were first subtracted using tracking beads to mark the flow-field and OpenPIV software. The ‘Gravity Machine’ analysis suite was used to determine sedimentation rate as described previously in Krishnamurthy et al. A high frequency filter was used for comparison of separate tracks. Raw traces vs. filtered data is presented in supplementary materials.

### Cell density sorter

The design was modified from that of Norouzi et al[34]. A design was modeled in Fusion360 and printed on a FormLabs 3+ instrument using Clearv4 resin. The channels were rinsed with isopropylalcohol before curing for 30 minutes. The channels were then flushed with liquid PTFE, then washed overnight in deionized water and .1% NP-40 for passivisation. The experiment was run using 2 syringe pumps set to accurately pass sample at .2ml/min. The output, or retrieval syringe, was filled with 1mL of F/2 medium at density 1.028. At the end of the sort, both syringes were emptied into a 50mm dish, and extra 1.028 density seawater was added so cells would sediment to the bottom of the dish for scoring.

### Light sheet imaging

Light sheet imaging of Pyrocystis cells was done on a Luxendo InVi SPIM instrument using a Nikon CFI Apo 25x W 1.1 NA objective with optivar set at either 31.3 or 62.5X total magnification. The sample was maintained in a water bath held at 21°C for the duration of each experiment. Image processing was done in Imaris. Isosurfaces were created for the fluorescence signal within the network and volumes were subsequently calculated. Since the cell wall has no fluorescent signal, another isosurface was projected on the exterior of the network encompassing the entire cell. The total calculated volume of this surface was used as a proxy of total cell volume. The Hexasphere in Fig. 3 was generated in Fusion 360.

### Long-term imaging

Long-term imaging of the Pyrocystis life-cycle was done on a custom ‘Octopi’ imaging system within a peltier controlled chamber kept at 21°C. The cells were imaged in a 50mm Pelco glass bottomed dish that was sealed with a glass coverslip after filling the chamber with F/2 medium. A LED panel was used to simulate the 12/12 light dark cycle.

### Electron Microscopy

Electron microscopy was done at the UC Berkeley electron microscopy core facility. Cell morphology was compared between samples fixed chemically and by high pressure freezing. Rapid chemical fixation provided the most reproducible results with minimal damage to overall morphology. Fixation was done with addition of 2% Paraformadahyde, 3% Gluteraldahyde to the media. Use of the Pelco BioWave was essential to avoid retraction of the cytoplasmic network or plasmolysis. The Pelco was used at 150watts, switched on for 1 minute with vacuum off, 1 minute with vacuum on, switched off for 1 minute with vacuum on, and finally 1 minute switched on with vacuum engaged. Samples were then stained and processed into resin before sectioning.

### RNA preparation for Long-Read sequencing

Around 5,000 cells were collected in a 40µm cell strainer and washed with 10mls of sterile filtered F/2 culture medium. Cells were then quickly scraped off the filter with a metal spatula and placed directly into 1ml of Trizol reagent where they were further homogenized for 30 seconds with a plastic pestle attached to a power drill. RNA was then purified on a column according to the PureLink Micro Kit protocol. RNA quality was assessed with a BioAnalyzer run using both the ‘Plant RNA pico’ and ‘Eukaryote RNA pico’ protocols. The RNA peaks on the electropherogram were considered high quality and processed further for sequencing by PacBio.

### Protein structure prediction

The transcriptome assembled from the long-read sequencing was first aligned against all entries in the UniProt/Swiss-Prot database. High quality alignments were then loaded into Alphafold using ChimeraX, and figures were generated using the same.

## Supplementary information

### METHODS

#### Obtaining dinoflagellates

Pyrocystis cells were first obtained from a CTD rosette bottle triggered at a depth of 75m. When foul weather prohibited further casts, vertical plankton tows were done from 25-35m with a 20 micron mesh plankton net at a max speed of 1kt. Presence of Pyrocystis noctiluca in the sample was confirmed via imaging on a flow-through Planktoscope instrument. Cells were then isolated on a dissection microscope and placed in filtered underway water. 5 micron polystyrene beads were added to the media prior to loading into the sample stage to mark flow fields for later PIV correction.

#### Cell Culture

Pyrocystis noctiluca (CCMP732) were ordered from Bigelow Labs and expanded in F/2 media(-Si) prepared using artificial seawater (reef salts) at a salinity of 36%, specific gravity 1.028. Cells were cultured at 21ºC under cool white fluorescent light at 2500 lux. Media was refreshed every 2 weeks to maintain proper salinity. Cells were expanded every 6 weeks to minimize accumulation of cell debris. For seawater density increase, a solution of Optiprep/2X seawater was prepared and added as needed to create isotonic seawater of higher density. Density was verified in a refractometer instrument.

#### Cell density sorter

The design was modified from that of Norouzi et al[**?**]. A design was modeled in Fusion360 and printed on a FormLabs 3+ instrument using Clearv4 resin. The channels were rinsed with isopropylalcohol before curing for 30 minutes. The channels were then flushed with liquid PTFE, then washed overnight in deionized water and .1% NP-40 for passivisation. The experiment was run using 2 syringe pumps set to accurately pass sample at .2ml/min. The output, or retrieval syringe, was filled with 1mL of F/2 medium at density 1.028. At the end of the sort, both syringes were emptied into a 50mm dish, and extra 1.028 density seawater was added so cells would sediment to the bottom of the dish for scoring.

## SUPPLEMENTARY THEORY

### Topological implications of inflation of reticulated cytoplasm

Here we explore the constraints associated with inflation of reticulated cytoplasm (n-genus torous) mapped onto a surface of a vacuole (spherical geometry) from a topological perspective. As the cell vacuole (approximated as a sphere of radius *r*) inflates post cell-division, the reticulated cytoplasm on the surface of the vacuole rearranges and strains to satisfy the inflated geometry (Fig. 3D). Although the reticulated cytoplasm forms a highly complex network of linked topology with the vacuole (Fig. 3A), here we only explore the implications associated with required membrane necessary for such a fast inflation event.

In order to calculate the total increase in membrane (or length) of the reticulated network as the vacuole expands several fold, we make several geometrical simplifications. Firstly, via imaging, we confirm that the inflation does not dramatically change the topology of the reticulated network which expands to accommodate a larger vacuole structure. Secondly, although the inflation is not entirely spherical, we assume a purely spherical geometry for the vacuole since it does not have major implications to a total estimate of membrane required for this inflation event. Lastly, instead of considering the complex reticulated geometry of the cytoplasm, we primarily consider an example using spherical tilings for this rough estimate.

As a vacuole expands in volume, total surface area of the sphere of radius *r* scales as 4*πr*^2^. Thus a vacuole embedded completely inside another spherical cytoplasm would require total membrane that scales with the square of the radius of the cell (*r*^2^). Here we demonstrate that for a reticulated mesh-like geometry (as is present in Pyrocystis) on the surface of a sphere, the required membrane during inflation scales linearly with the size of the cell (*r*).

For simplicity, we will consider a hexasphere tiling where majority of the surface of the sphere is covered with a hexagonal grid, with 12 pentagonal tiles. This tiling can be easily derived by sequential bifurcation along the edges of an icosahedral followed by a popping event. The original 12 vertices of the icosohederal harbor the 12 pentagons while the rest of the entire surface of the sphere can be tiled with a hexagonal grid. Let’s consider the tiled hexagon represents the reticulated network on the surface of the vacuole - with edge length of the hexagon represented with length *l*. Since area of a regular n-gon is 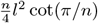, the area of the flat plane captured by the hexagon on a hexasphere is 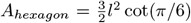 . As we assume that the topology of the reticulated network does not change with the inflation event, the total area of the sphere tiled this way is composed of *m* hexagons and 12 pentagons. For *m >>* 12, we can approximate the surface area of the sphere with *m*.*A*_*hexagon*_ ignoring the area covered by the 12 pentagons.

Rearranging, length of the hexagon on a hexasphere can be expressed in terms of the area terms as 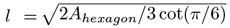. Substituting *A*_*hexagon*_ ≈*A*_*sphere*_*/m* where we ignore the area of 12 pentagons (for *m >>* 12). Thus total length of the network comprising a hexasphere can be expressed as *L ≈* 3*ml*. Substituting *l*, we can estimate 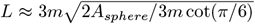or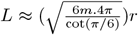. As can be seen, the length of the hexasphere grid scales linearly with increase in radius of the sphere *r*. Thus to first approximation, increase in the vacuole size as the cell inflates only requires membrane in proportion to the vacuole radius as compared to square of the radius in case of a vacuole that is completely enveloped by a cytoplasm. This reticulated architecture thus enables a large cellular expansion without requiring a large amount of membrane. As was previously demonstrated in Fig. 3g&h, this extra membrane comes from membrane folds visible in TEM cross sections.

### Gravitational Trap

Gravity as a fundamental force of nature has likely shaped the evolution of marine phytoplankton. If they succumb to the downwards gravitational pull, i.e. if the net buoyancy is negative at all times, the suspended cells will eventually leak to the abyss, thus rendering many marine life-forms extinct. Au contraire, if the net buoyancy is always positive they would remain afloat in the nutrient dearth environment, again leading to the same fate. Evidently, phytoplankton can both rise and sink. Thus a question naturally arises – how do their vertical trajectories in the ocean look like? Here, we present a minimal theoretical framework within which to couple gravitational sinking and an ecosystem profile to cell “behavior”, thus placing ecosystem dynamics of sinking/rising plankton in an ambit of driven Stokesian suspensions.

In the regime of low Reynolds numbers and large Peclet numbers, the dynamics is governed by the steady Stokes equation. Consider a spherical cell of radius *r* containing fluid of density *ρ*_*cell*_, and suspended in an ambient sea water with density *ρ*_*sw*_ and dynamic viscosity *µ*. The vertical velocity of the cell is given by [1]

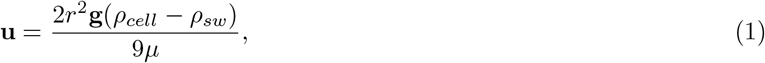

where the gravity direction **g** is aligned with the - z-axis and homogeneity and isotropy is assumed in the horizontal *x−y* plane. We divide the parameters in the above equations into two classes – external (environmental) *ρ*_*sw*_ and internal (behavioral) *r, ρ*_*cell*_ physical degrees of freedom, and the physics is introduced through the Stokes’ law (1). Activity manifests in this system through a time dependent internal state of the cell, which is lumped into one nondimensional variable *ϕ* = *αt/nT*_*c*_ where *T*_*c*_ is the time period of the cell cycle and *n* = 1, 2, 3.. is the number of cell cycles, and all the internal physical degrees of freedom depends on *ϕ*. Note that we have introduced the activity through the parameter *α*, which drives the cell cycle at a constant rate *α/T*_*c*_. The central idea is that as *ϕ* changes, cells can oscillate in the vertical spatial direction. Taking into account the compartmentalization of fluids within the cell, the vertical velocity can be written as

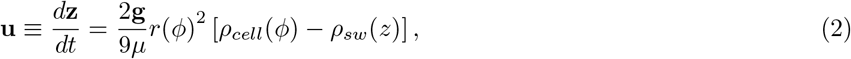

where *ρ*_*cell*_(*ϕ*) = *ρ*_*cy*_*V*_*cy*_*/V*_*c*_ + *ρ*_*vac*_*V*_*vac*_*/V*_*c*_ is the effective density of the cell which is a function of the internal state *ϕ* of the cell, with the *V*_*cy*_, *V*_*vac*_, *V*_*c*_ are the volume of cytoplasm, vacuole and the cell respectively. Note that these expressions are independent of the precise geometry of the cytoplasm, and in fact vacuole and cytoplasm can have non-trivial geometry; and the sea water density *ρ*_*sw*_ is a function of depth *z*.

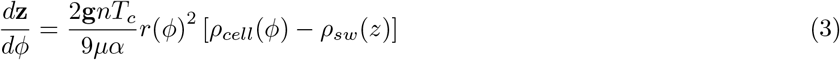

First, we explore the above equation in the ballooning phase, where the *V*_*vac*_ increases by bringing in fresh water from the ambient sea water, via selective permeability.

Non-dimensionalizing lengths by *L* = *r*_*min*_ and times by *T* = 9*µ/*(2*g*(*ρ*_*cy*_ *− ρ*_*vac*_)*r*_*min*_) gives

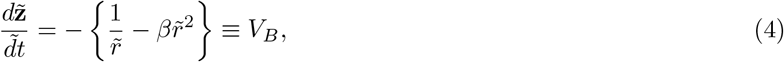

where 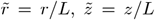 and 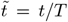 and we have introduced a non-dimensional density dependent parameter *β*(*z*) = (*ρ*_*sw*_(*z*) *− ρ*_*vac*_)*/*(*ρ*_*cy*_ *− ρ*_*vac*_). The above equation has a stable fixed point

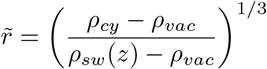

at which cells become neutrally buoyant. Assuming only fresh water permeates through the cell membrane filling the vacuole *ρ*_*vac*_ = 1000*kg/m*^3^, and using the CTD data from Kilo Moana as a representative density profile in the ocean, we get *ρ*_*sw*_ = 1024.3 + 0.9 tan^−1^ 0.05(*z−* 107.8)*kg/m*^3^, and from the gravity machine data we get *ρ*_*cy*_ = 1040*kg/m*^3^. These typical values during ballooning gives a phase portrait shown in Fig. S7. To further analyse the above equations, we turn to the experiments for the time dependence of radius during ballooning which displays saturation behavior for *ϕ»*1. This motivates an inverse tangent relationship between the cell radius *r* and cell phase *ϕ* in the ballooning phase

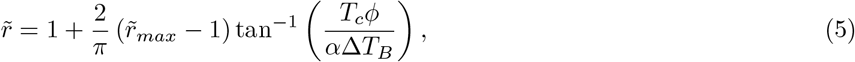

where 1*/α*Δ*T*_*B*_ = 0.0032, *r*_*min*_ = 48.77*µm* and *r*_*max*_ *r*_*min*_ = 18.6*µm*. And its derivative w.r.t. *ϕ* becomes a Lorentzian function,

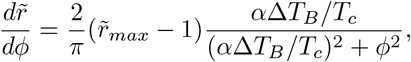

with a width 2*α*Δ*T*_*B*_*/T*_*c*_ and maximum 0.38*T*_*c*_*/α*Δ*T*_*B*_. With these ingredients, equation (4) can be solved for 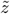in terms of 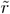, away from the regions of sharp gradient in the pycnocline where *β*(*z*) can be assumed to be constant.

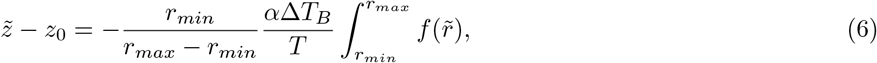

where

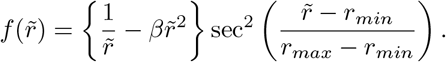

The full dynamical behavior of the cell, however, requires a priori knowledge of *ρ*_*cell*_(*ϕ*), throughout the cell cycle. We approach this problem with a simplifying assumption of considering two stages in the cell cycle that consist of –1) the fast times of rapid expansion Δ*T*_*B*_*/T*_*c*_ during which the cell density changes sharply, and 2) slow times of cell growth Δ*T*_*A*_*/T*_*c*_ in which the density grows gradually.

The form of equation (6) has a stretched coordinate Φ = *ϕ/δ*, where *δ*≡*α*Δ*T*_*B*_*/T*_*c*_ *«*1, naturally forming a boundary layer [2] of thickness *δ* in the temporal density profile at *ϕ* = 0. From equation (4) the density in the inner region where Φ = 𝒪 (1), is given by

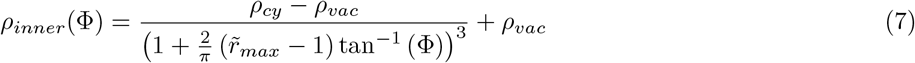

Assuming linear increase in effective cell density during the growth phase after inflation, the density in the outer region *ϕ » δ*, to leading order in *δ* takes the form

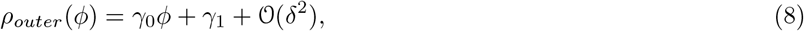

and the asymptotic matching of the inner and outer densities at the edge of the boundary layer

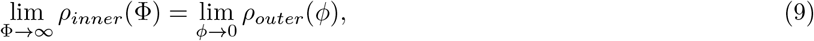

gives the intercept 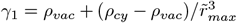in (8), and the constraint of periodicity on all cell cycle parameters gives the slope *γ*_0_ = (*ρ*_*cy*_*−γ*_1_)*/*(1*−δ*); ensuring that the effective cell density returns back to its initial state at *ϕ* = 0. (7) and (8) gives the effective density that is valid in the full domain of *ϕ* of the cell cycle

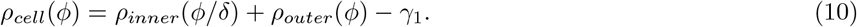

With these ingredients, the dynamical equation in the 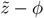 space becomes

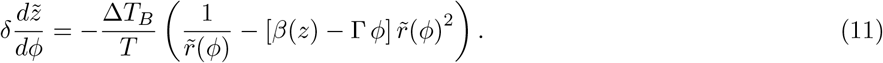

Here Δ*T*_*B*_*/T* is the ratio of ballooning time scale to sedimentation time scale, 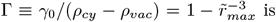 is the measure of relative density reduction due to ballooning.

Note that a small parameter *δ* multiplies with the derivative in equation (11), and therefore ballooning event can be seen as a singular perturbation to the dynamics [2], where *δ* = *α*Δ*T*_*B*_*/T*_*c*_ is the boundary layer thickness in *ρ*_*cell*_ created due to the abrupt ballooning event, and *β*(*z*) = (*ρ*_*sw*_(*z*) *− ρ*_*vac*_)*/*(*ρ*_*cy*_*−ρ*_*vac*_) captures the pycnocline. Also, note that although the mathematical forms of *ρ*_*inner*_ and *ρ*_*outer*_ are motivated by experimental observations specific to a given species, the matched asymptotic approach presented above can be applied to richer class of cell cycle phases, that may involve multiple boundary layers of various thicknesses in time domain.

Equation (11) is amenable to singular perturbation analysis in which the leading order solution is just the solution to equation (4). However, the lifetime of the ballooning event *δ* is negligibly small compared to the total transit time of the organism, thus only the outer solution of equation (11) is ecologically relevant. In particular, the ecological and mathematical relevance of the ballooning event comes in resetting the slingshot and adding residual corrections to the trajectory as a perturbation expansion in *δ*. We get the outer solution to (11) by replacing 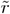 with 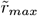and integrating with respect to *ϕ* from *δ* to *α*, and retaining terms to leading order in *δ*.

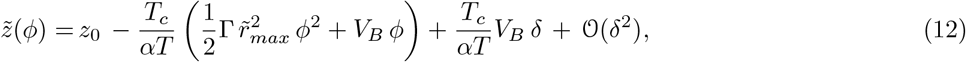

where *V*_*B*_ is the velocity immediately after ballooning, given by (4) with 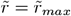 [see Fig. 2 E], and *z*_0_ is the ocean depth at which the first cell cycle begins. (12) is a parabola with a vertex at 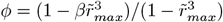.

From this we can see that the critical ballooning radius required for escaping the gravity trap, *r*_*critical*_ = (*β −*0.5)^−1*/*3^ diverges as *β→* 0.5 with a critical exponent 1*/*3.

Note that, when the pycnocline is constant, equation (11) can be mapped, to leading order in *δ*, to the equation of a projectile shot vertically

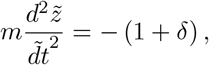

with an effective mass or inertia 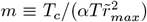 and effective drive 1 + *α*Δ*T*_*B*_*/T*_*c*_ along negative z axis, both of which are dependent on the activity *α* in the system; where we have defined activity as a force that drives the cell cycle clock in Fig. 3.

For a complete solution of (11) we carry out numerical integration and find escaping and trapped trajectories in Fig. 5, for small and large values of *ρ*_*cy*_ respectively.

### Aquaporin Number-density Estimation

Volume of the pre-inflated cell:

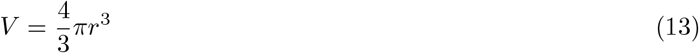

Substituting the value of *r* of a pre-inflated cell:

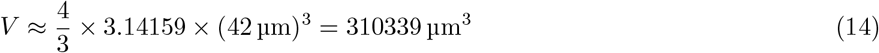

convert the volume from cubic micrometers to nanoliters:

Pre inflated volume = .3103*nL*

Substituting the value of *r* of a post-inflated cell:

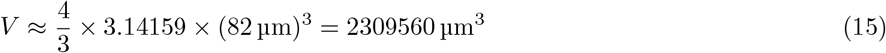

convert the volume from cubic micrometers to nanoliters:

Post inflated volume = 2.31*nL*

Flux=**2nl**

6.66 *×* 10^16^ molecules of water

To calculate the surface area of a pre-inflated cell:

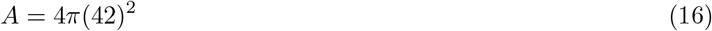

Pre inflated surface area = 22167*µm*^2^

To calculate the surface area of a post-inflated cell:

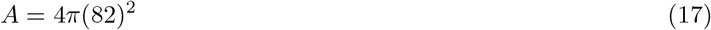

Post inflated surface area = 84496*µm*^2^

### Fitting inflation

The log-logistic function can be written as:

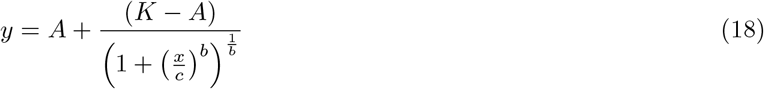

Fit for experimental data of radius increase:

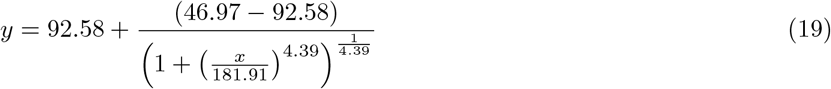

Target water transport =6.66 *×* 10^16^molecules

Estimated transport capacity = 1*×*10^9^molecules per second per channel

Timescale = 1000 seconds

Minimal number of channels needed =2.4*/µm*^2^ at start, .7*/µm*^2^ at end

**FIG. S1.**
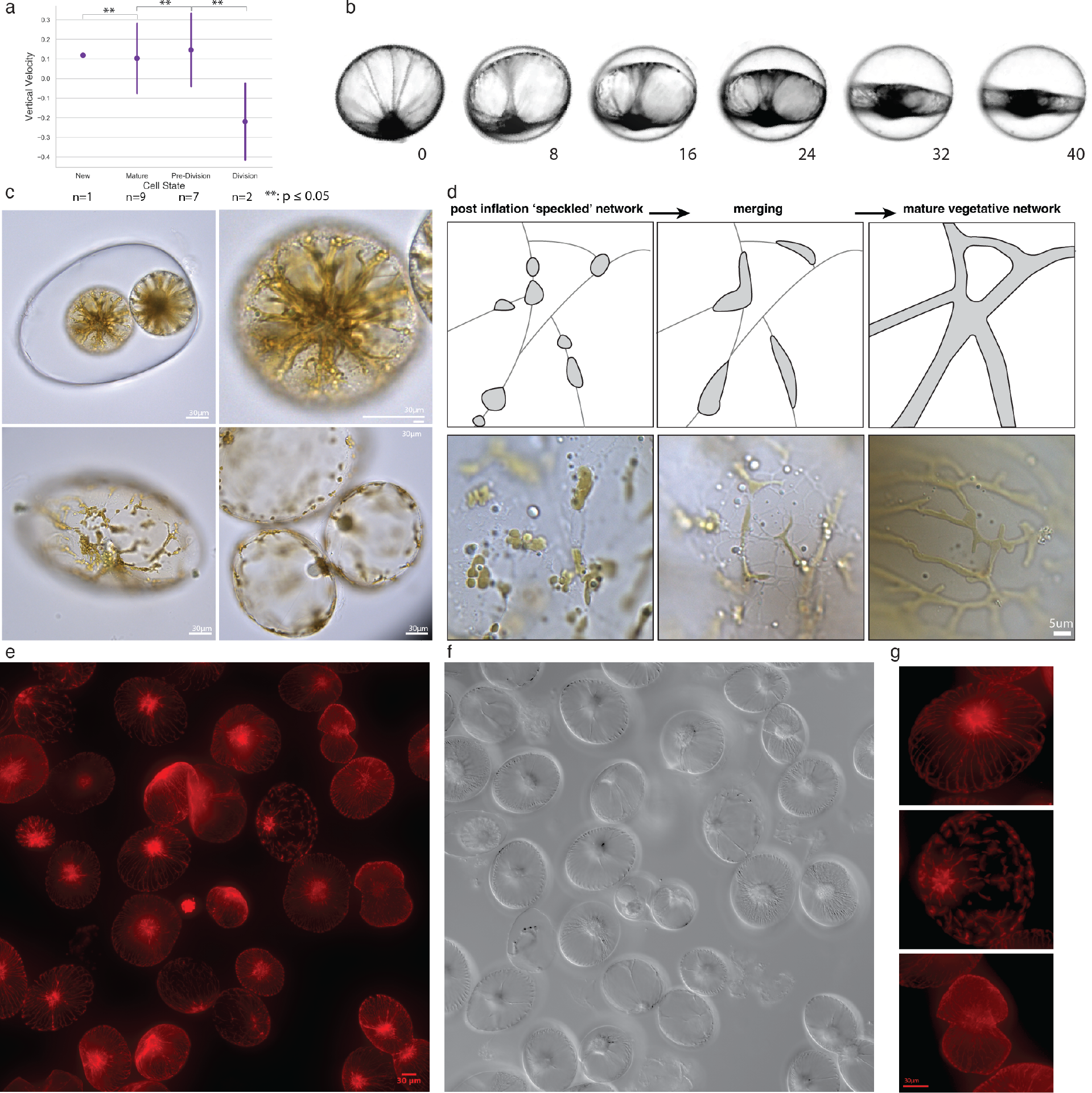
A)Average velocity values for the various Pyrocystis cell states obtained on board HOT317. Error bars represent standard deviation, and p-values from pairwise t-tests indicate the statistical significance of the differences in average velocities between states B)Preceding division, Pyrocystis undergoes large changes in the volumetric ratios between intercellular (apoplastic) and vacuolar space, while largely keeping the external hydrodynamic profile the same. C) Pyrocystis has a cytoplasmic network that is largely condensed in the pre-inflated daughter cells. As the cell inflates, the cytoplamic contents appear ‘speckled’ throughout the extended network. D) In the hours after inflation, the speckled cytoplasmic contents begin to merge, forming the mature vegetative network. E) The plastids within the cytoplasmic network display strong autofluorescent signal under red excitation channels. F)Transmitted channel from D. G)The cell states are displayed in the various cytoplamic configurations.

**FIG. S2.**
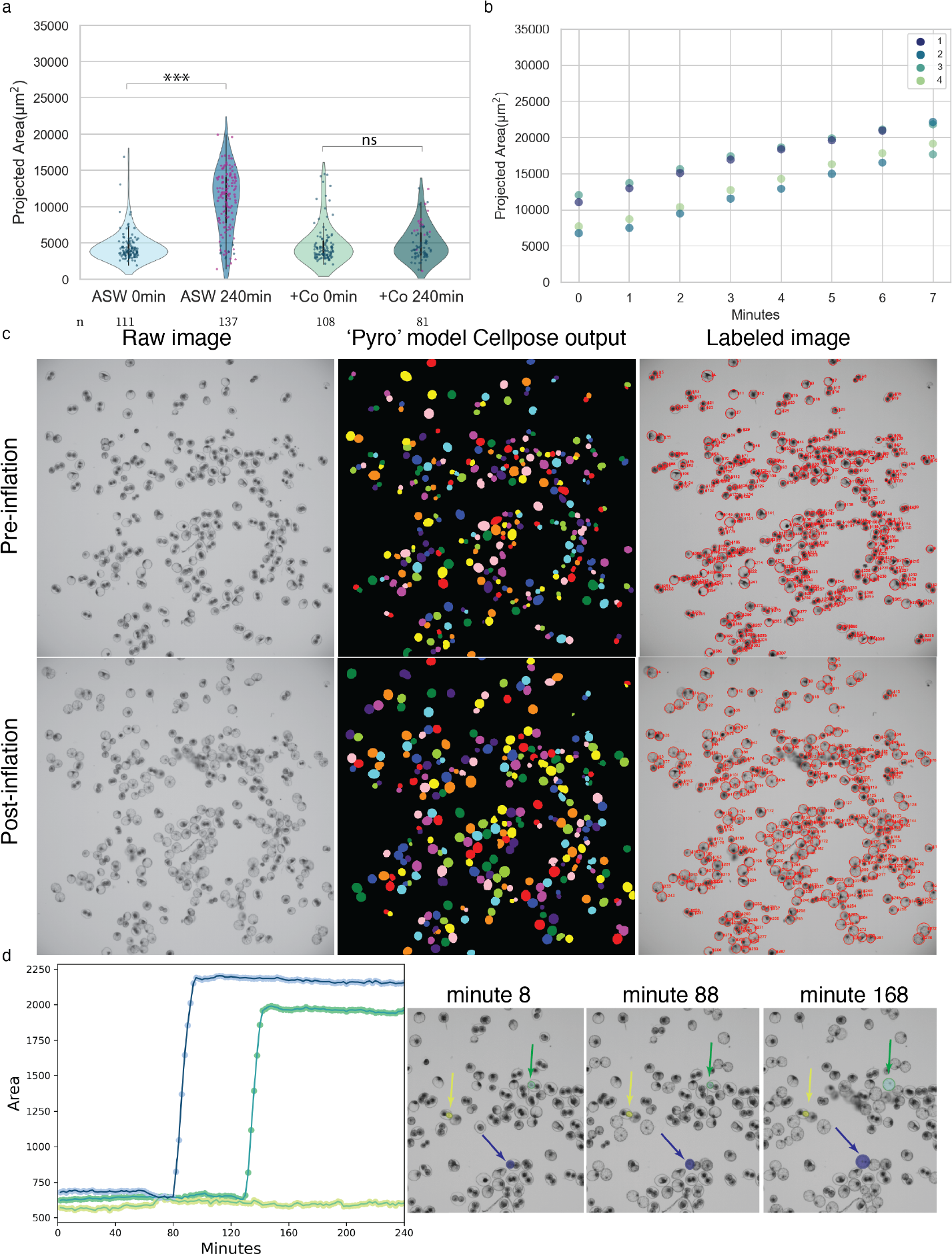
A)Ion-channel blockers such as Cobalt also inhibit inflation in Pyrocystis. B)Premature rupturing of the cell wall in dividing cells induces expansion of daughter cells. Figure is quantification of cells in Movie S11. The mother cell wall surrounding cells 1 and 2 were ruptured 2 minutes prior to cells 3 and 4. C) Segmentation pipeline for projected area calculations. Cell masks were generated by re-training the Cyto2 model in Cellpose using roughly 300 pre-division cells. A centroid tracking script tracked individual cells and their projected areas through consecutive time points.D)Three representative cells are tracked through the time series.

**FIG. S3.**
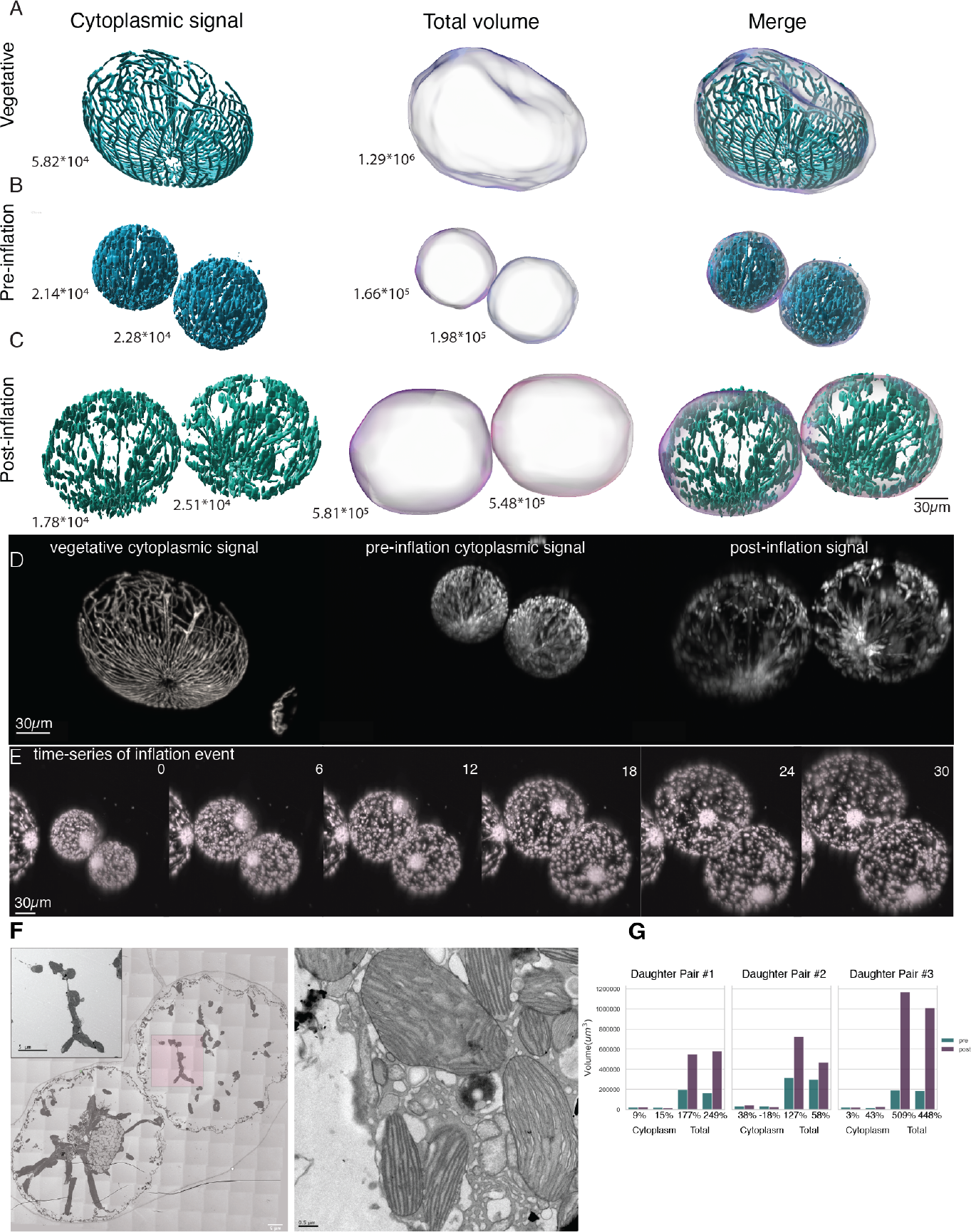
A)The fluorescence signal from the cytoplasm was used to generate a volume surface within Imaris, from which content volume was calculated. An isosurface was generated around the cytoplasmic signal to estimate total cell volume .B)In a pre-division cell, the cytoplasmic signal is condensed in a much smaller total cell volume. C)After 15 minutes of inflation, the cytoplasm extends without increasing substantially in volume. D)Raw fluorescence signal used for volumetric estimations.E) Fluorescence signal from an inflating pair of daughter cells immediately following cell wall breakdown.F)The cytoplasmic strands are compressed in pre-division cells. These strands in Pyrocytis are all highly jammed with numerous organelles and vacuoles. G)Quantification of cytoplasmic vs. total cell volume increase during the process of inflation for 3 pairs of Pyrocystis division events.

**FIG. S4.**
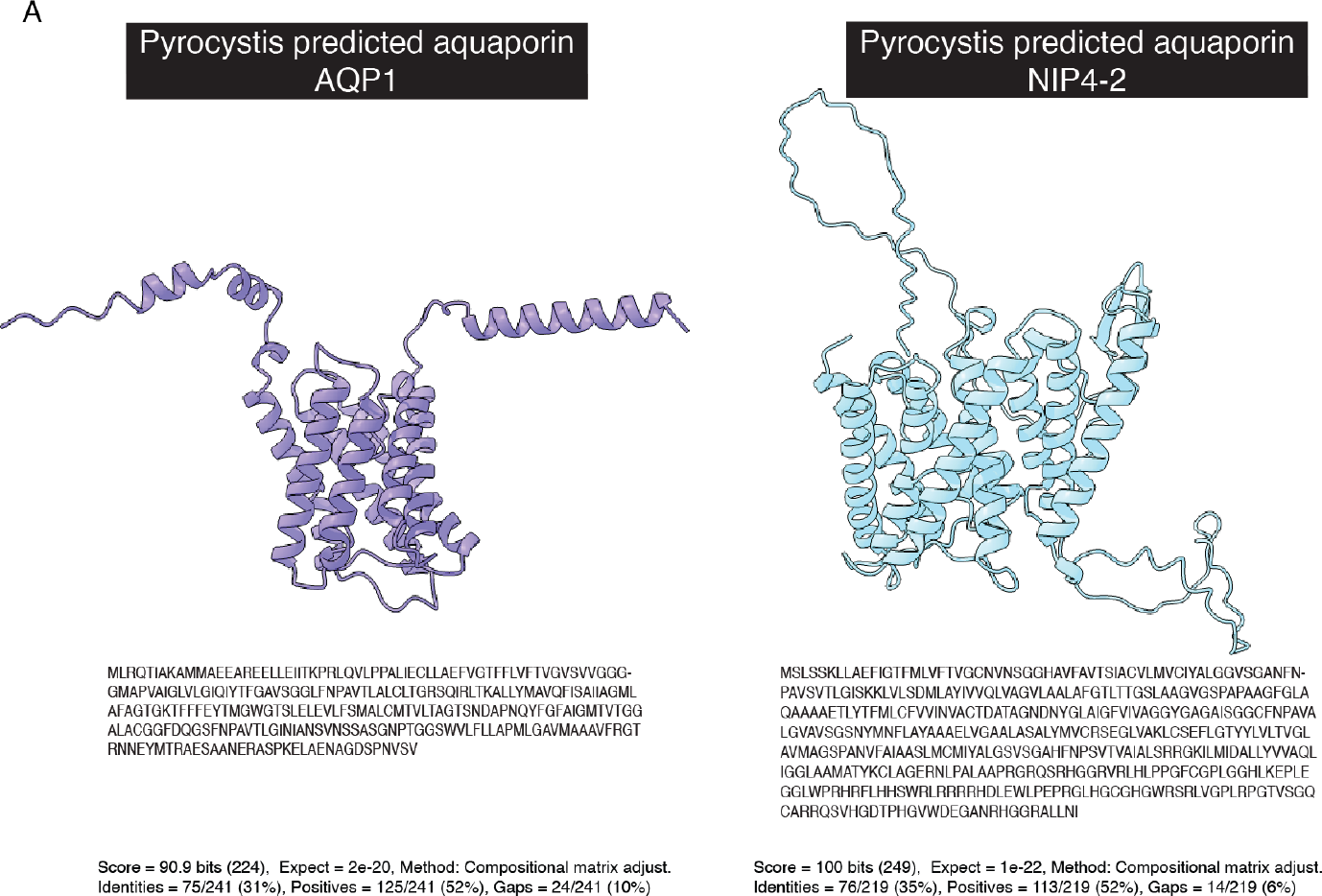
A)Sequence alignments using transcriptomic assemblies from long-read sequencing reveal proteins related to existing aquaporin channel proteins. Structure prediction from Alphafold reveals configurations consistant with transmembrane channels.

**FIG. S5.**
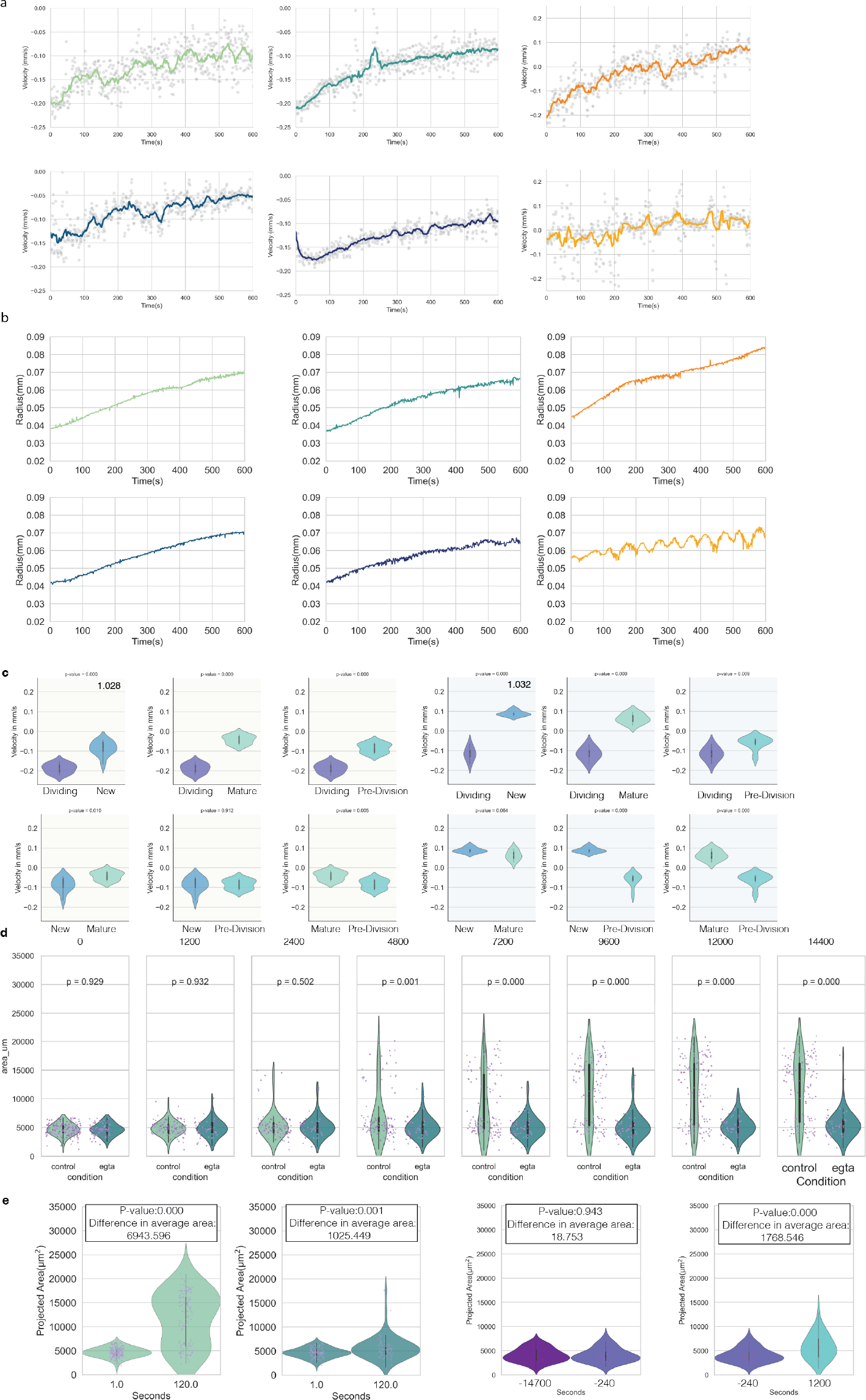
A)Raw velocity plots from 6 different cell division events within the vertical tracking microscope with denoised data overlayed. Blue/green hues are 1.028 seawater density, orange hues are 1.032. B) Radius vs. Time plots from the 4 different cell division events reported above. C) Results of t-test comparing weighted velocity measurements from Fig. 2. D) Results of t-test comparing control and EGTA treated timepoints from Fig. 2. E) Results of t-test comparing area change in Fig. 2. Violin plots show the distribution of the projected area (µm2) over time for different conditions. The width of the violin indicates the density of data points at different values of projected area. The white dot represents the median (50th percentile), the thick black line indicates the interquartile range (IQR; 25th-75th percentiles), and the thinner black lines show the lower and upper adjacent values. The overlaid strip plot displays individual data points, jittered along the x-axis for better visibility.

**FIG. S6.**
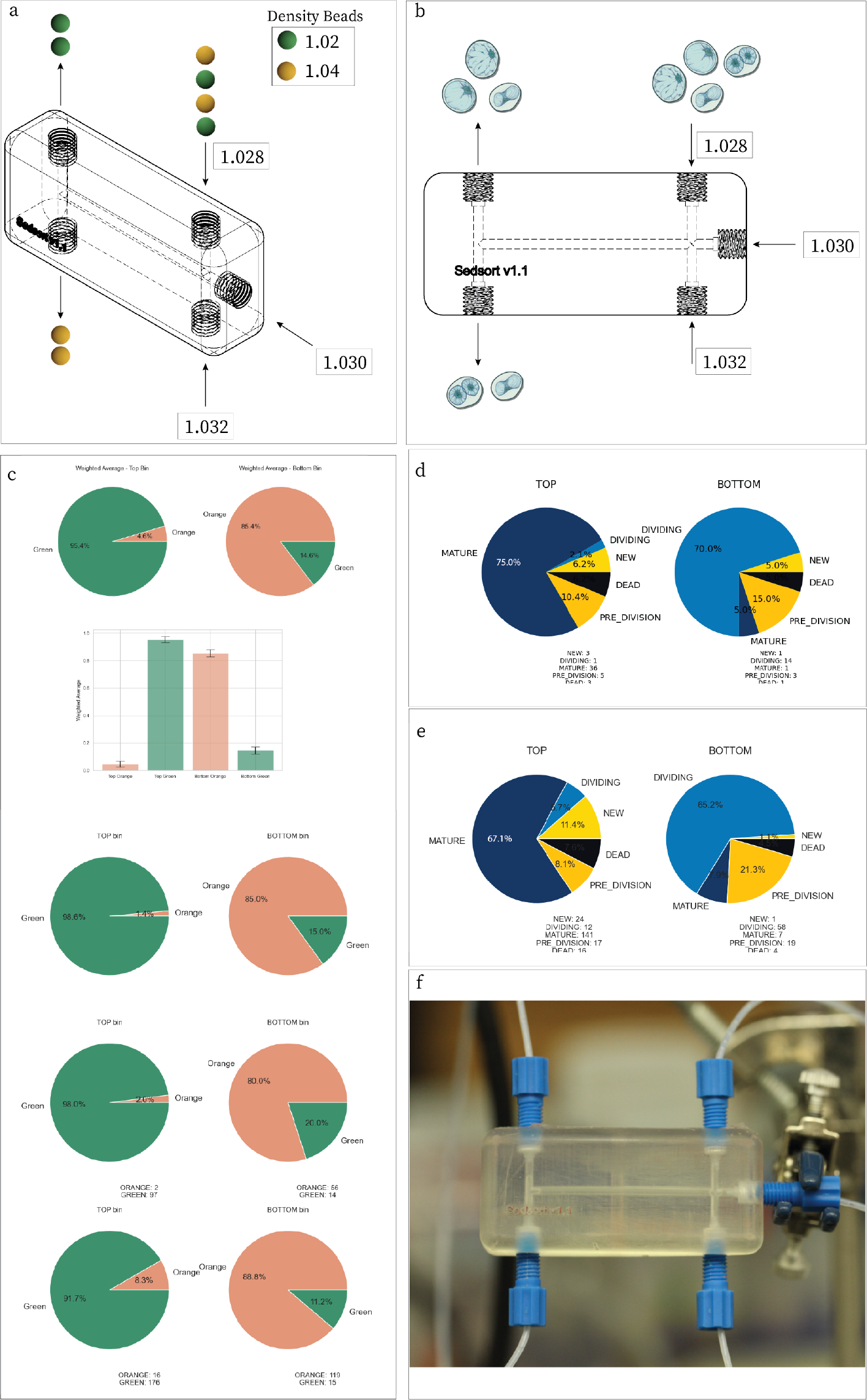
A)A 3D printed sedimentation based cell sorter was validated with density beads in seawater of varying densities. B) The sorter bins Pyrocystis cells based on cell state, agreeing with density measurements calculated from sedimentation velocity values. C) Quantification of device validation shown in A. D) Quantification of experiment shown in B, flowing around 100 cells through the sorter E) Results of flowing a well-mixed large culture through the sorter. F) Image of the experimental setup.

**FIG. S7.**
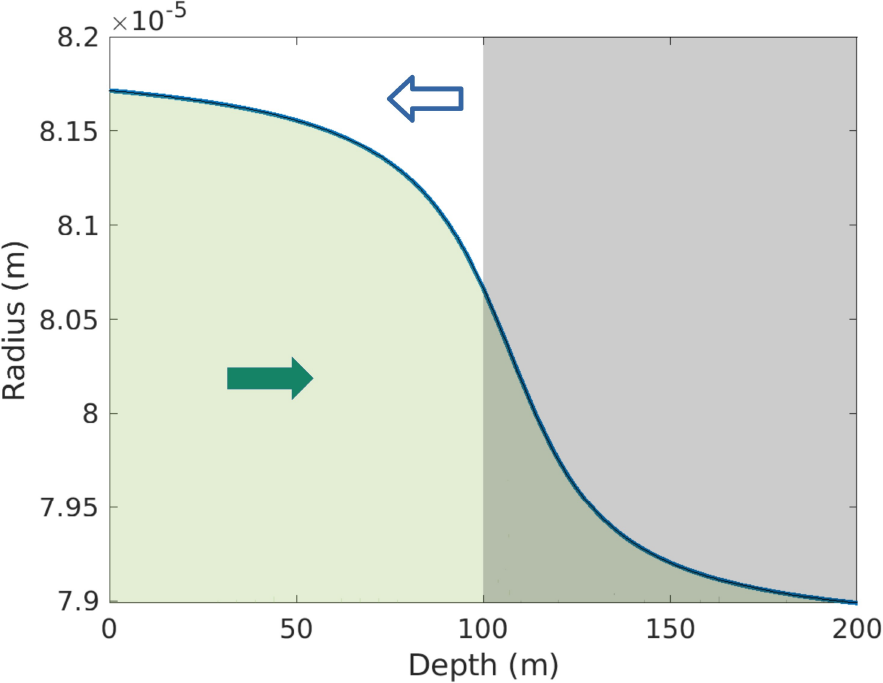
Sinking and rising in the ballooning phase

**FIG. S8.**
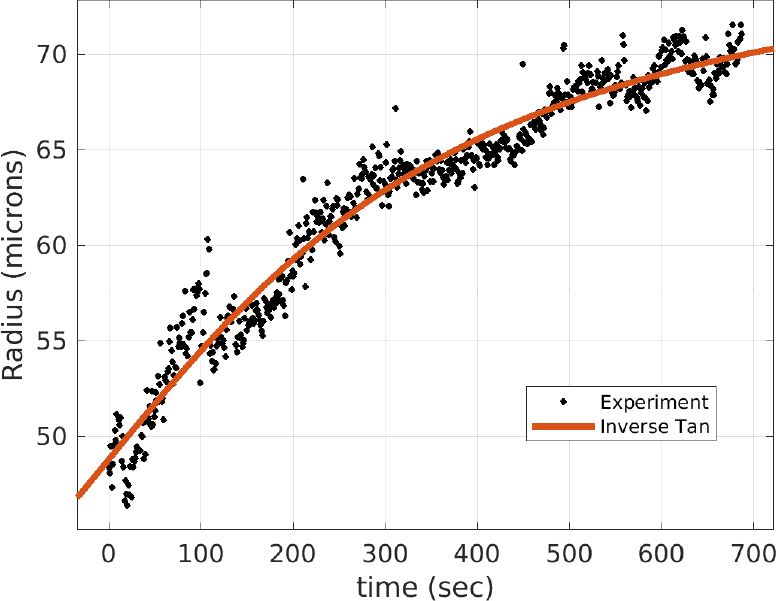
Ballooning radius

**FIG. S9.**
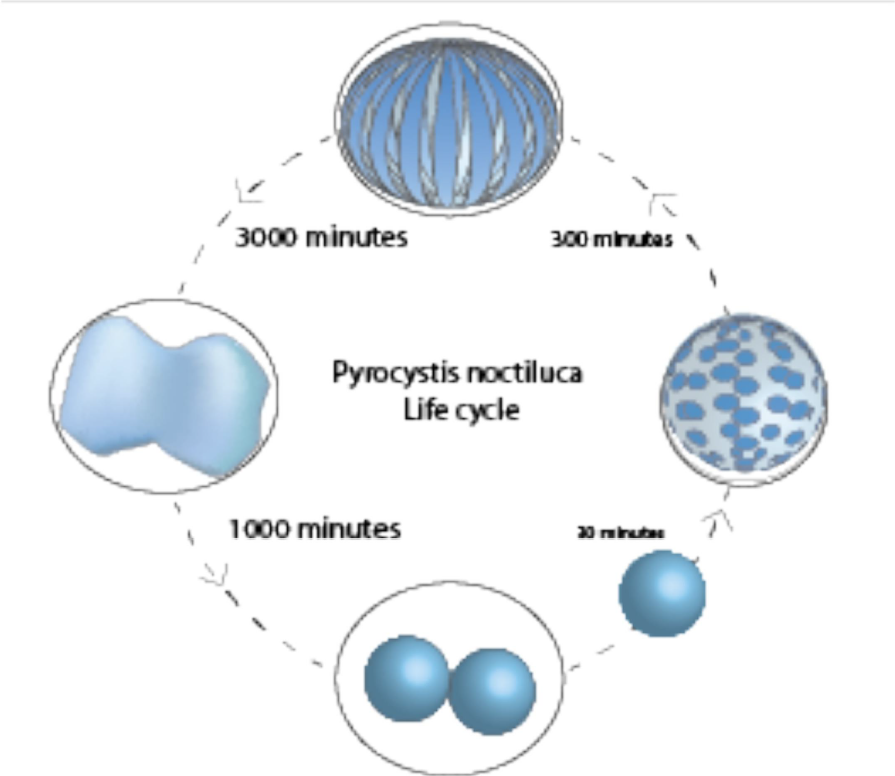
Internal states of the cell

**FIG. S10.**
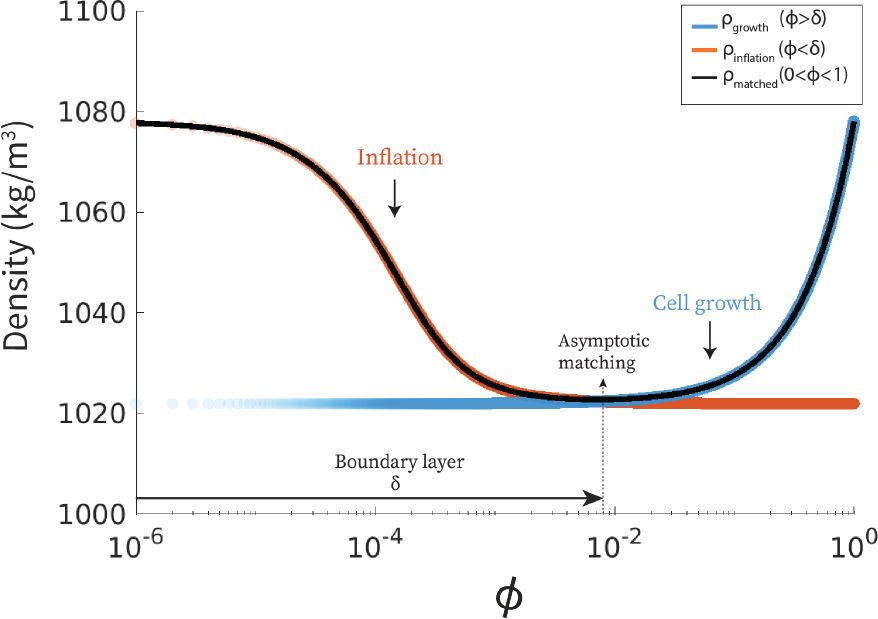
Matching inner and outer density at the boundary layer for 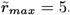.

**FIG. S11.**
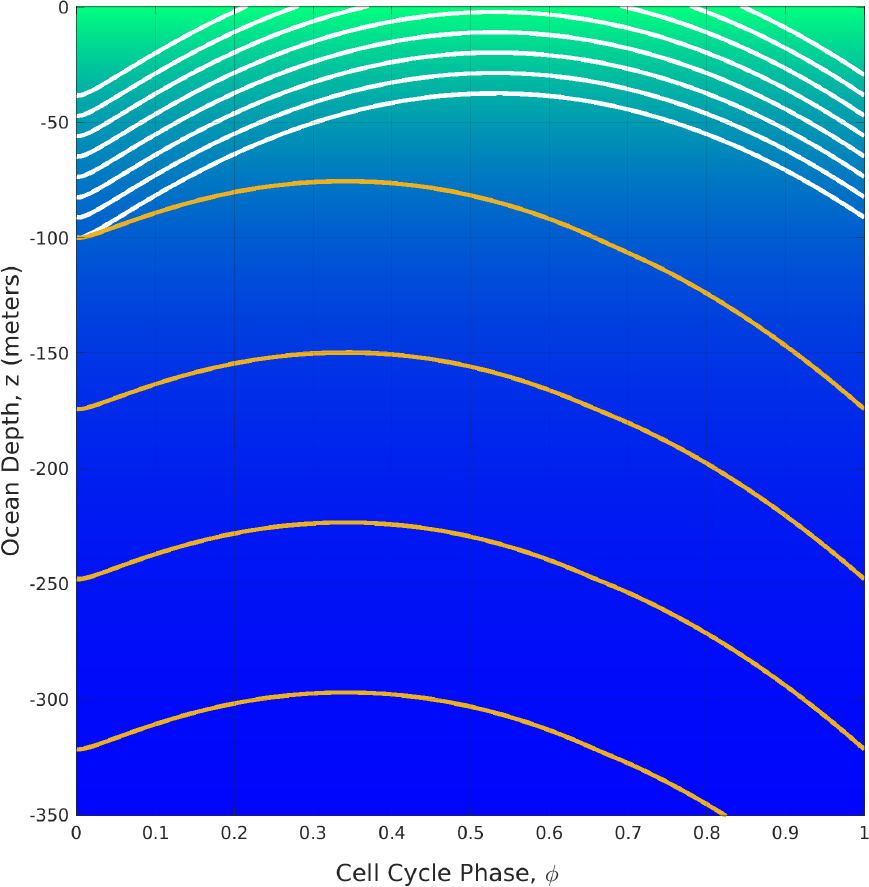
Cell succumbing to and escaping from the gravity trap for activity.

**FIG. S12.**
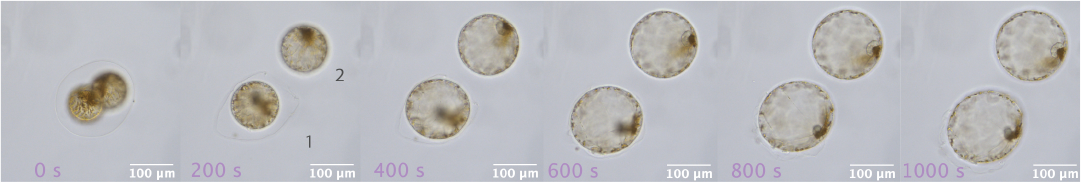
Post-division inflation.

**FIG. S13.**
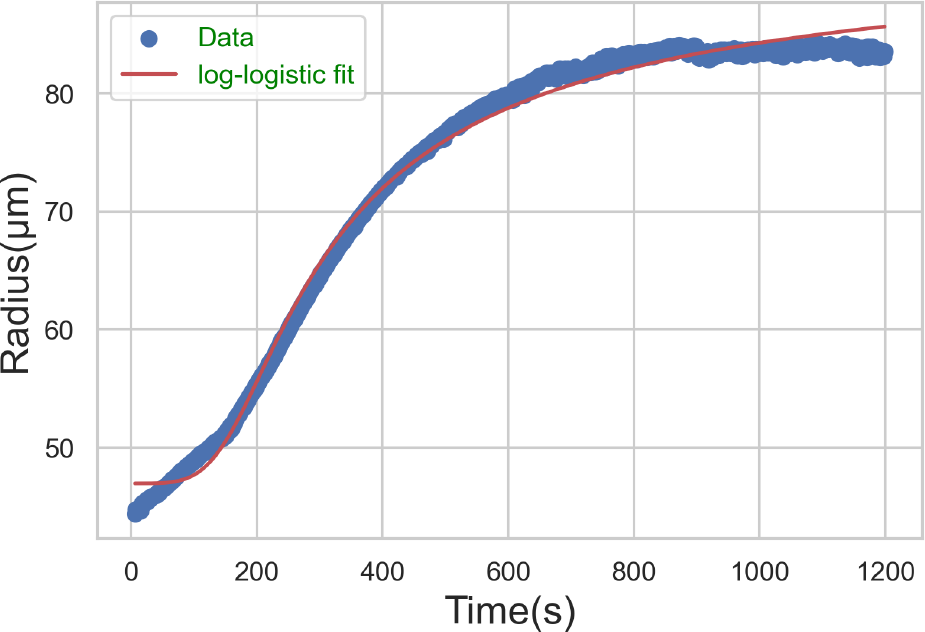
log-log fit of experimental inflation data from time-lapse microscopy.

**FIG. S14.**
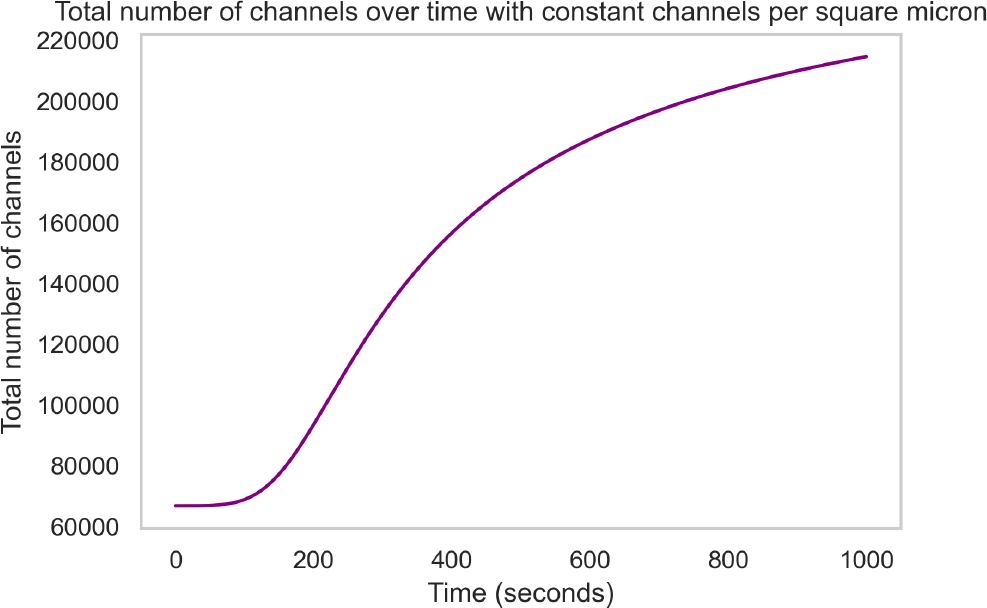
Estimation of total number of aquaporin channels on surface, given estimated conditional parameters for water flux.

